# Impact of competition between precursor and mature microRNAs on stochastic gene expression

**DOI:** 10.64898/2026.01.03.697472

**Authors:** Raunak Adhikary, Dipjyoti Das

## Abstract

MicroRNAs (miRNAs) are key post-transcriptional regulators, processed from precursor miRNAs (pre-miRNAs) into mature miRNAs through nuclear and cytoplasmic proteins. Recent evidence shows that pre-miRNAs and mature miRNAs can compete for the same target mRNAs, yet the impact of this miRNA maturation-driven competition on gene expression noise remains unknown. We address this in widespread feedback motifs where both pre- and mature miRNAs degrade a protein’s transcripts, and the protein itself either activates or represses miRNA transcription. Using a mathematical model, we show that miRNA maturation tunes the behavior of positive or negative feedback loops, which function as bistable switches or oscillators at the mean-field level, respectively. The relative degradation of mature versus pre-miRNAs and the mRNA-miRNA co-degradation rates can jointly modulate the parameter regions of bistability or oscillations. Moreover, for positive feedback, stochastic simulations reveal that bimodal mRNA distributions emerge near the saddle-node bifurcation boundaries, but not always within the bistable regions. Bimodal mRNA distributions also appear for negative feedback, but outside the region of limit cycles. Importantly, in both feedback types, such noise-induced bimodality emerges in regions where mean-field analysis predicts no bistability or limit cycles. These results demonstrate that noise-induced phenotypic variability cannot necessarily be linked to underlying deterministic bifurcations and elucidate how miRNA maturation shapes stochastic gene expression in regulatory motifs relevant to development and disease.

## 1 Introduction

Cells tune transcription and translation in response to environmental cues [103]. However, these processes are inherently noisy due to small numbers of molecular players and fluctuating environmental conditions. This noise drives striking cell-to-cell variation in mRNA and protein copy numbers, even among genetically identical cells — a phenomenon termed as ‘gene expression noise’ [50, 52, 82, 85]. This variability in gene expression can contribute to cell fate change and survival in fluctuating environments [7, 55, 69, 97]. Deciphering how cells regulate this noise has become a significant theme in molecular biology [25, 74, 81, 88, 96].

RNA polymerases (RNAPs) initiate transcription by binding to promoter DNA, while transcription factors (TFs) binding to nearby sites modulate this process by enhancing or repressing RNAP activity [85]. Both theoretical [21, 24, 86, 87, 89, 91] and experimental studies [18, 29, 39, 48, 80, 95] have shown that stochastic TF binding/unbinding shapes the promoter architecture and plays a key role in transcriptional regulation of gene expression noise. In addition, TF-mediated positive feedback on transcription was found to drive bimodal protein distributions. For example, Choi et al. used the *lac* operon system in *E. coli* to show that the LacY gene forms an autoregulatory positive feedback and variation in inducer concentration (which inhibits the *lac* repressor) leads to bimodal LacY expression [18]. Also, Zander et al. used the natural lactose uptake system of *E. coli*, which consists of the same *lac* operon, to show bimodality in LacZ expression [115].

Although transcriptional regulation of gene expression noise is well studied, its post-transcriptional control by short noncoding RNAs remains less understood. In prokaryotes, small RNAs (sRNAs) bind and degrade target mRNAs [36–38, 54, 60, 72], while in eukaryotes, various types of small RNAs down-regulate gene expression. Among them, microRNAs (miRNAs) are of notable interest for their roles in stress response, development, neuronal function, and cancer metastasis [14, 42, 57, 62, 64, 67, 92].

RNA polymerase II molecules transcribe miRNA genes into primary transcripts (typically ¿ 1 kb), which are cleaved in the nucleus by proteins like Drosha and DGCR8 to generate precursor miRNAs (pre-miRNAs, 60–90 nt) [23, 58, 59]. The pre-miRNAs are then exported to the cytoplasm, where Dicer and associated proteins further process them into mature miRNAs ( ∼ 22 nt) [68, 109]. Stress-induced DNA damage and certain tumor conditions can disrupt this maturation process [71, 102], indicating that precise regulation of miRNA maturation is essential for normal cellular function.

Mature miRNAs typically bind to the 3^*′*^ untranslated region (UTR) of target mRNAs. The nucleotide position 2-7 of the 5^*′*^ end of miRNA, known as the ‘miRNA seed’, is vital in target recognition and remains conserved under strong selection pressure [4, 19, 40, 104]. Since this seed sequence remains conserved during miRNA maturation, precursor miRNAs can, in principle, interact with the same target mRNAs and compete with their mature counterparts [17]. For instance, the primary miRNA pri-let-7 can repress *lin*-41 mRNA even in the absence of processed mature let-7 [99, 113]. Similarly, pre-miR-124 competes with mature miR-124 for 3^*′*^ UTR binding sites on *SNAI2* mRNA [84], and pre-miR-21 shares overlapping binding sites with mature miR-21 in the 3^*′*^ UTR of *TGFBR2*, influencing its translation [15].

Experiments have shown that tuning miRNA concentration can establish a threshold in target mRNA expression, separating highly repressed and expressed regimes [73]. Theoretical studies have investigated this miRNA-mediated threshold behavior using different modeling frameworks. Some models assumed a fixed pool of miRNAs [73], while others explicitly modeled the miRNA transcription of a miRNA-coding gene and its interaction with target mRNAs [76]. Furthermore, some models [9, 10] explored the non-linear cross-talks among multiple target mRNAs that share a common miRNA pool - a phenomenon known as the ‘competing endogenous RNAs (ceRNA)’ [3, 30, 94]. These studies predicted that stochastic fluctuations in ceRNA networks can generate bimodal mRNA distributions, which was later supported by experimental observations [10, 22].

Transcription factor (TF)–mediated and miRNA-mediated regulations often act together in gene regulatory networks controlling key physiological and developmental processes, including cell cycle progression [12, 26, 111], cell fate determination [43, 77, 105], cancer proliferation [19, 27, 116], angiogenesis [6, 16, 31, 121], and host–HIV interactions [32]. Feedback and feedforward loops involving miRNAs are widespread in such networks [47, 70, 100, 110]. Among them, miRNA-mediated positive [11, 28, 49, 61, 79, 101, 107, 118] and negative feedback loops [53, 70, 100, 112, 119] are particularly common and represent the simplest yet fundamental network motifs.

Most theoretical studies on miRNA-mediated feedback employ nonlinear deterministic models under the mean-field approximation. Such models have shown that miRNA-mediated positive feedback can generate bistability or multistability [2, 13, 41, 65, 66, 98], whereas negative feedback can produce oscillatory dynamics in the mean target expression [35, 106, 108, 120]. Some theoretical studies focused on building stochastic frameworks to investigate the noise in target gene expression and revealed that miRNAs can buffer the noise [8, 63, 75, 93, 98, 117]. It was also predicted that miRNA-based negative feedback can lead to noise-induced bimodal distributions of target mRNAs near the cross-over between the expressed and repressed regimes [1, 76]. However, existing models overlook that precursor and mature miRNAs can compete for a shared pool of target mRNAs. How this competition, mediated by the miRNA maturation process, shapes noise and stability in gene expression remains unexplored.

Here, we develop a stochastic model that explicitly incorporates miRNA maturation and captures competition between precursor and mature miRNAs for a shared target mRNA pool. Depending on whether proteins translated from the target mRNAs activate or repress the miRNA gene, the resulting miRNA-mediated feedback is negative or positive, respectively. Mean-field analysis reveals that the steady-state mean mRNA shows bistability with saddle-node bifurcations in the positive feedback, and oscillations with Hopf bifurcations in the negative feedback, in accordance with earlier studies [56, 120]. Crucially, the relative degradation rates of precursor and mature miRNAs, along with their efficiencies in degrading target mRNAs, significantly influence the extent of bistability and oscillatory regimes. Stochastic simulations further reveal the emergence of bimodal mRNA distributions, driven by enhanced intrinsic noise near the saddle-node bifurcation boundaries in the positive feedback loop. Notably, this bimodality can persist outside the deterministic bistable region. Similarly, in negative feedback, we observe noise-induced bimodality, similar to earlier models lacking the miRNA maturation process [1, 76]. This bimodality arises solely from noise amplification and is confined to regions outside the limit-cycle regime. Thus, in both feedback architectures, stochasticity generates bimodal expression in parameter regions where mean-field theory predicts neither bistability nor oscillations. Together, these results yield testable predictions showing that competition between precursor and mature miRNAs significantly alters the parameter space supporting bimodal gene expression compared to models without such competition.

## 2 Model

In this study, we extend a previously published model of miRNA-mediated feedback [120] by incorporating miRNA maturation to examine the competition between precursor and mature miRNAs for a common target mRNA pool. In the model, the mRNAs are synthesized from a gene at a rate *k*_*r*_ and translated into proteins at a rate *k*_*p*_. The mRNAs and proteins degrade with rates *g*_*r*_ and *g*_*p*_, respectively. Precursor miRNAs (pre-miRNAs) are synthesized from the miRNA coding gene at a basal rate 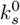 and converted to mature miRNAs at a rate *k*_*mat*_ via the maturation process. The precursor and mature miRNAs degrade with rates *g*_*s*_ and *g*_*m*_, respectively. We assume that both pre- and mature miRNAs bind irreversibly to target mRNAs, co-degrading each other at respective rates *γ*_*s*_ and *γ*_*m*_, consistent with the stoichiometric interaction between miRNAs and mRNAs [5, 51, 78]. Moreover, the proteins act as transcription factors (TFs) that bind to the miRNA gene at a rate *k*_*b*_ and unbind at a rate *k*_*u*_, switching the gene between ‘TF-bound’ and ‘TF-unbound’ states. In the TF-bound state, pre-miRNAs are synthesized at a rate *k*_*s*_. Notably, when 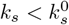, the TF binding establishes positive feedback on the target gene expression; when 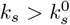, the feedback is negative. We characterize the feedback strength by the dimensionless ratio *β* = *k*_*b*_*/k*_*u*_.

The model can further be described by the following set of reactions:

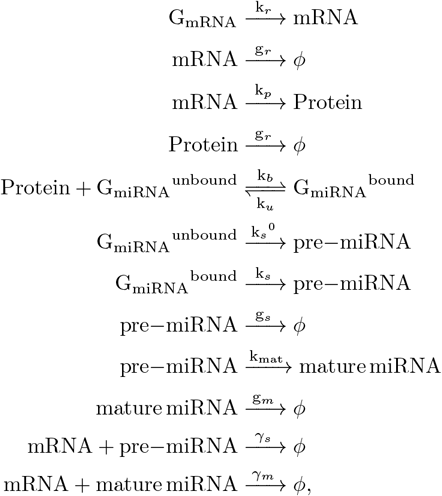

Where 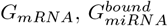, and 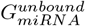 denote the mRNA-coding gene, the miRNA-coding gene in ‘TF-bound’ and ‘unbound’ states, respectively.

Based on the reaction scheme, we can write down the chemical Master equation describing the joint probability distribution of mRNA, protein, pre-miRNA, and mature miRNA copy numbers, by *r*(*t*), *p*(*t*), *s*(*t*), and *m*(*t*), respectively (see SI, section 1). In principle, moment equations for the means and variances can be derived from the Master equation. However, due to its strong nonlinearity, these equations form an infinite hierarchy that cannot be closed without approximations [20, 90]. To obtain analytical insights, we derived approximate equations for the means (see SI, section 3). In addition, we performed exact stochastic simulations using the Gillespie algorithm [33, 34], with kinetic parameters taken from previous studies (Table 1).

**Table 1.** Description of parameters and their values.

| Symbol | Description | Value | References |
| --- | --- | --- | --- |
| $k_r$ | mRNA synthesis rate | varied ($0.0s^{-1} - 0.6s^{-1}$) | [1, 9] |
| $g_r$ | mRNA degradation rate | $0.0004s^{-1}$ | [1, 44] |
| $k_p$ | protein synthesis rate | $0.017s^{-1}$ | [120] |
| $g_p$ | protein degradation rate | $0.00002s^{-1}$ | [1, 44] |
| $k_b$ | protein binding rate of miRNA-coding gene | varied ($10^{-7}s^{-1} - 10s^{-1}$) | [1] |
| $k_u$ | protein unbinding rate of miRNA-coding gene | $10^{-2}s^{-1}$ | [1] |
| $k_s^0$ | pre-miRNA synthesis rate in ‘unbound’ state | $\begin{cases} 0.5s^{-1} & \text{for positive feedback} \\ 0.05s^{-1} & \text{for negative feedback} \end{cases}$ | [1, 9] |
| $k_s$ | pre-miRNA synthesis rate in ‘bound’ state | $\begin{cases} 0.05s^{-1} & \text{for positive feedback} \\ 0.5s^{-1} & \text{for negative feedback} \end{cases}$ | [1, 9] |
| $g_s$ | pre-miRNA degradation rate | varied ($0.000017s^{-1} - 0.00017s^{-1}$) | [83, 122] |
| $k_{mat}$ | pre-miRNA maturation rate | varied ($10^{-7}s^{-1} - 10s^{-1}$) | Assumed in this study |
| $g_m$ | mature miRNA degradation rate | varied ($0.000017s^{-1} - 0.00017s^{-1}$) | [83, 122] |
| $\gamma_s$ | mRNA-pre-miRNA co-degradation rate | varied ($0.004s^{-1} - 4s^{-1}$) | Assumed in this study |
| $\gamma_m$ | mRNA-mature-miRNA co-degradation rate | varied ($0.004s^{-1} - 4s^{-1}$) | Assumed in this study |

Under the mean-field approximation (i.e., neglecting fluctuations), we derived the equations of mean mRNA (⟨*r*⟩), protein (⟨*p*⟩), pre-miRNA (⟨*s*⟩), and mature miRNA (⟨*m*⟩) from the Master equation as given below (see SI, Eqs. S3.1–S3.5):

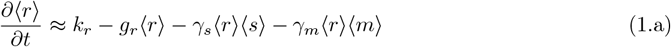

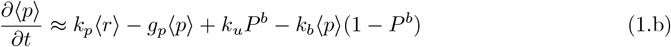

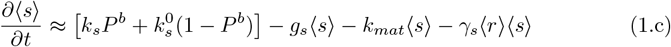

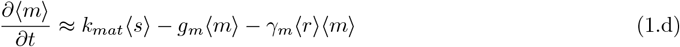

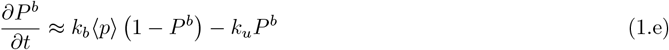

Here, e, ⟨·⟩ denotes the ensemble average, and *P* ^*b*^ is the probability of the miRNA-coding gene to be in the ‘bound’ state. In the steady state, the mean mRNA is given by the following polynomial equation.

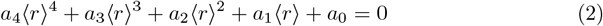

where

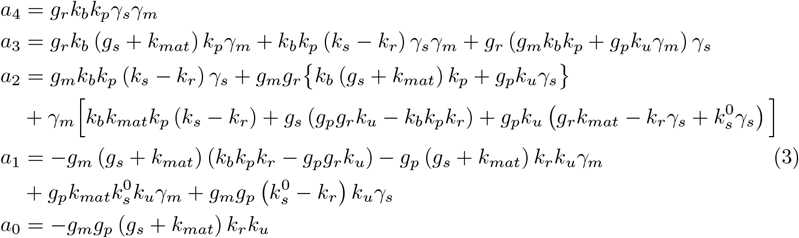

We solve Eq. 2 numerically for different parameters and perform stability analysis of the steady-state solutions (see sections 4, 5, and 6 in the SI). Below, we summarize the results of mean-field calculations and exact stochastic simulations.

## 3 Results

### 3.1 Impact of miRNA Maturation on the Target Expression in a Positive Feedback Loop

#### 3.1.1 miRNA maturation can modulate the extent of bistable regimes

We first focus on the positive feedback loop (i.e.,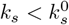) to explore the roles of pre-miRNA maturation rate (*k*_*mat*_) and miRNA degradation rates (*g*_*s*_ and *g*_*m*_) in modulating the mean expression in the steady state. In Fig. 2, we show the mean mRNA as a function of mRNA transcription rate (*k*_*r*_) for different combinations of *k*_*mat*_, *g*_*s*_, and *g*_*m*_. Analysis of our mean-field equations (Eqs. 1.a-1.e) shows that the mean mRNA exhibits bistability, as already reported in earlier studies without miRNA maturation [120]. The red vertical lines in Fig. 2 represent the saddle-node bifurcation points on the *k*_*r*_-axis (denoted by 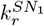 and 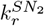), obtained from the stability analysis of the mean-field equations (see SI section 5 and 6). A region of bistability exists between these bifurcation points.

To validate the mean-field results, we performed exact stochastic simulations with two distinct initial conditions: zero and high nonzero initial copy numbers of mRNAs, proteins, pre- and mature miRNAs ((*r*_0_, *p*_0_, *s*_0_, *m*_0_ = 0, 0, 0, 0) and (*r*_0_, *p*_0_, *s*_0_, *m*_0_ = 2000, 2000, 2000, 2000)). We find that the mean mRNA ( ⟨*r*⟩) from stochastic simulations agrees well with deterministic calculations (Fig. 2). The bifurcation points can capture the transitions from low to high values of ⟨*r*⟩ for both zero and nonzero initial conditions. Note that, in Fig. 2, the left bifurcation point 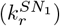 at a lower *k*_*r*_ value corresponds to the nonzero initial condition, whereas the right bifurcation point 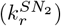 at a higher *k*_*r*_ value corresponds to the zero initial condition. This is because high initial numbers of mRNAs and proteins keep the miRNA gene mostly in the bound state, resulting in lower miRNA synthesis rate due to positive feedback. Thus, for high nonzero initial condition, the transition from high to low values of ⟨*r*⟩ occurs at a lower mRNA transcription rate (*k*_*r*_), as compared to the case of zero initial condition.

Notably, when mature miRNAs have longer lifetimes than pre-miRNAs (i.e., *g*_*s*_ *> g*_*m*_), the region of bistability widens with increasing pre-miRNA maturation rate (*k*_*mat*_) (Fig. 2A, B). Conversely, when pre-miRNAs have longer lifetimes than mature miRNAs (*g*_*s*_ *< g*_*m*_), the bistable region shortens with increasing *k*_*mat*_ (see Fig. 2C, D). This change in the bistable region occurs due to the shift of the right bifurcation point 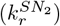 with varying *k*_*mat*_, but the left bifurcation point 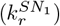 remains the same (Fig. 2). To understand this, we first note that the mean mRNA mainly depends on the individual degradation rates of pre- and mature miRNAs since the co-degradation rates of ‘mRNAs-pre-miRNA’ and ‘mRNA-mature-mRNA’ pairs are kept same for Fig. 2 (i.e., *γ*_*s*_ = *γ*_*m*_). When mature miRNAs are more stable than pre-miRNAs (*g*_*s*_ *> g*_*m*_), the mRNAs are degraded more by the mature miRNAs with increasing *k*_*mat*_ that ensures faster conversion of pre-miRNAs into mature miRNAs (see the model in Fig. 1). Thus, when we start from zero initial condition, as *k*_*mat*_ increases, ⟨*r*⟩ remains low until a higher mRNA transcription rate (*k*_*r*_) is reached, pushing the transition point 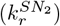 to a higher *k*_*r*_ value (comapre Fig. 2A and 2B). On the other hand, if pre-miRNAs are more stable than mature miRNAs (*g*_*s*_ *< g*_*m*_), mRNAs are more targeted by relatively unstable mature-miRNAs as *k*_*mat*_ increases. Therefore, ⟨*r*⟩ switches from a low to a high value at a lower *k*_*r*_, shifting the transition point 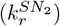 leftward with increasing *k*_*mat*_ (Fig. 2C, D). This demonstrates how the parameters, *k*_*mat*_, *g*_*s*_, and *g*_*m*_ modulate the bistable region.

**Figure 1.**
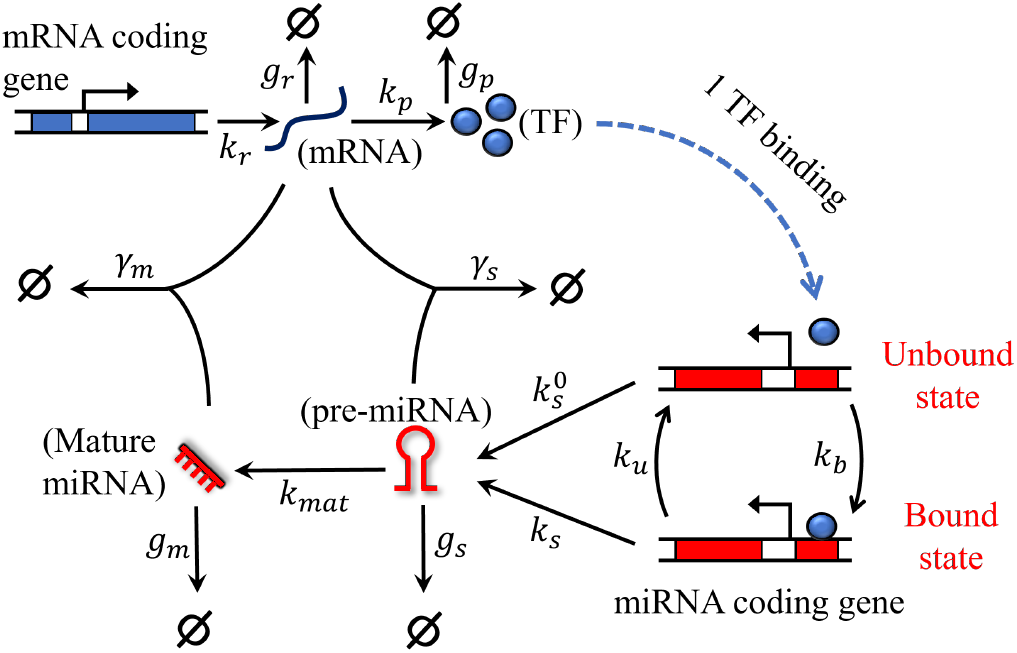
Schematic model showing the miRNA maturation process and miRNA-mediated feedback loop. Various kinetic processes and corresponding rates are shown by arrows (see the Model section and the parameters in Table 1). The precursor miRNAs (pre-miRNAs) and mature miRNAs interact with the target mRNAs, and both “pre-miRNA-mRNA” and “mature-miRNA-mRNA” pairs are mutually codegraded. Free mRNAs produce proteins, which act as transcription factors (TF) and affect the pre-miRNA transcription rate. The miRNA-coding gene toggles between TF-bound and TF-unbound states, in which the miRNA transcription rates are *k*_*s*_ and 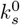, respectively. Depending on whether 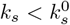 or 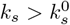, the TF binding establishes positive or negative feedback, respectively. Here, the ratio of TF binding to unbinding rates (*β* = *k*_*b*_*/k*_*u*_) can be considered as the feedback strength.

#### 3.1.2 miRNA degradation and mRNA–miRNA codegradation jointly tune bistability

We next explore the role of miRNA-mRNA co-degradation rates (*γ*_*s*_ and *γ*_*m*_) on the bistable expression in the parameter space of mRNA transcription rate (*k*_*r*_) and pre-miRNA maturation rate (*k*_*mat*_). In Fig. 3, we show the bistable regions demarcated by the bifurcation boundaries (solid red curves obtained from mean-field analysis) for various combinations of *γ*_*s*_, *γ*_*m*_, *g*_*s*_, and *g*_*m*_. Note that the bifurcation points in the *k*_*r*_-axis in Fig. 2 become bifurcation curves in the *k*_*r*_-*k*_*mat*_ plane in Fig. 3. To compare this with the stochastic simulations, we define ‘the absolute difference of steady-state mean mRNA’ as

**Figure 2.**
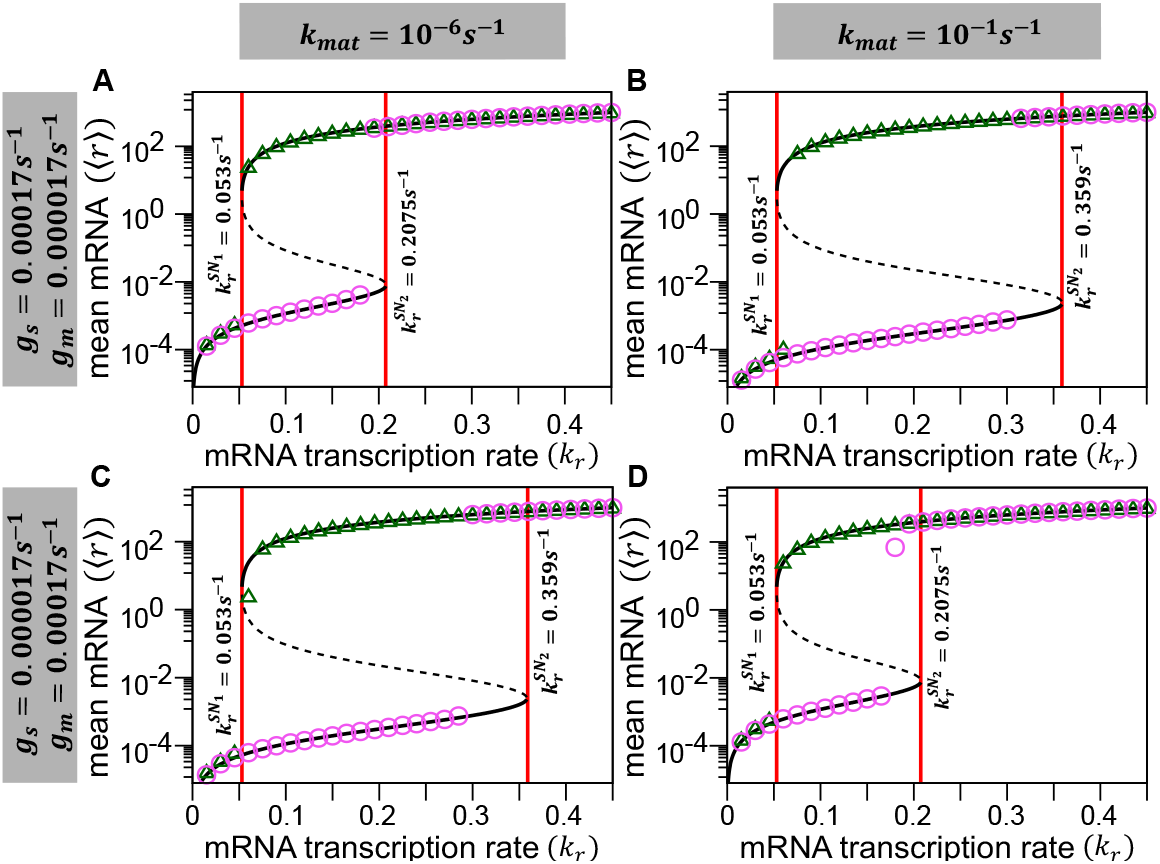
Bifurcation diagrams show that miRNA maturation modulates the bistability region in the positive feedback loop. The steady-state mean mRNA ( ⟨*r*⟩) is plotted against the mRNA transcription rate (*k*_*r*_) for different pre-miRNA maturation rates and for varying degradation rates of pre- and mature miRNAs. Black solid and dashed curves represent stable and unstable solutions of the mean-field equations, respectively (see SI, sections 5 and 6, for stability analysis). Red vertical lines denote the saddle-node bifurcation boundaries (denoted by 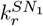 and 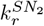). Magenta circles and green triangles represent steady-state ⟨*r*⟩ obtained from stochastic simulations for two different initial conditions, respectively: zero ((*r*_0_, *p*_0_, *s*_0_, *m*_0_) = (0, 0, 0, 0)) and high nonzero ((*r*_0_, *p*_0_, *s*_0_, *m*_0_) = (2000, 2000, 2000, 2000)) initial copy numbers of the mRNAs, proteins, pre-miRNAs and mature miRNAs. Top (A-B) and bottom panels (C-D) are obtained for two different conditions, namely when mature miRNAs are more stable and less stable than pre-miRNAs (*g*_*s*_ *> g*_*m*_ and *g*_*s*_ *< g*_*m*_, respectively, but *γ*_*s*_ = *γ*_*m*_). In A-B, the region of bistability widens as the pre-miRNA maturation rate (*k*_*mat*_) increases, while the bistable region shrinks with increasing *k*_*mat*_ in C-D. Parameters: *k*_*b*_ = 10^−3^*s*^−1^, *k*_*u*_ = 10^−2^*s*^−1^(i.e.,*β* = 0.1 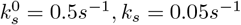, *γ*_*s*_ = *γ*_*m*_ = 0.04*s*^−1^, and other parameters are from Table 1.

**Figure 3.**
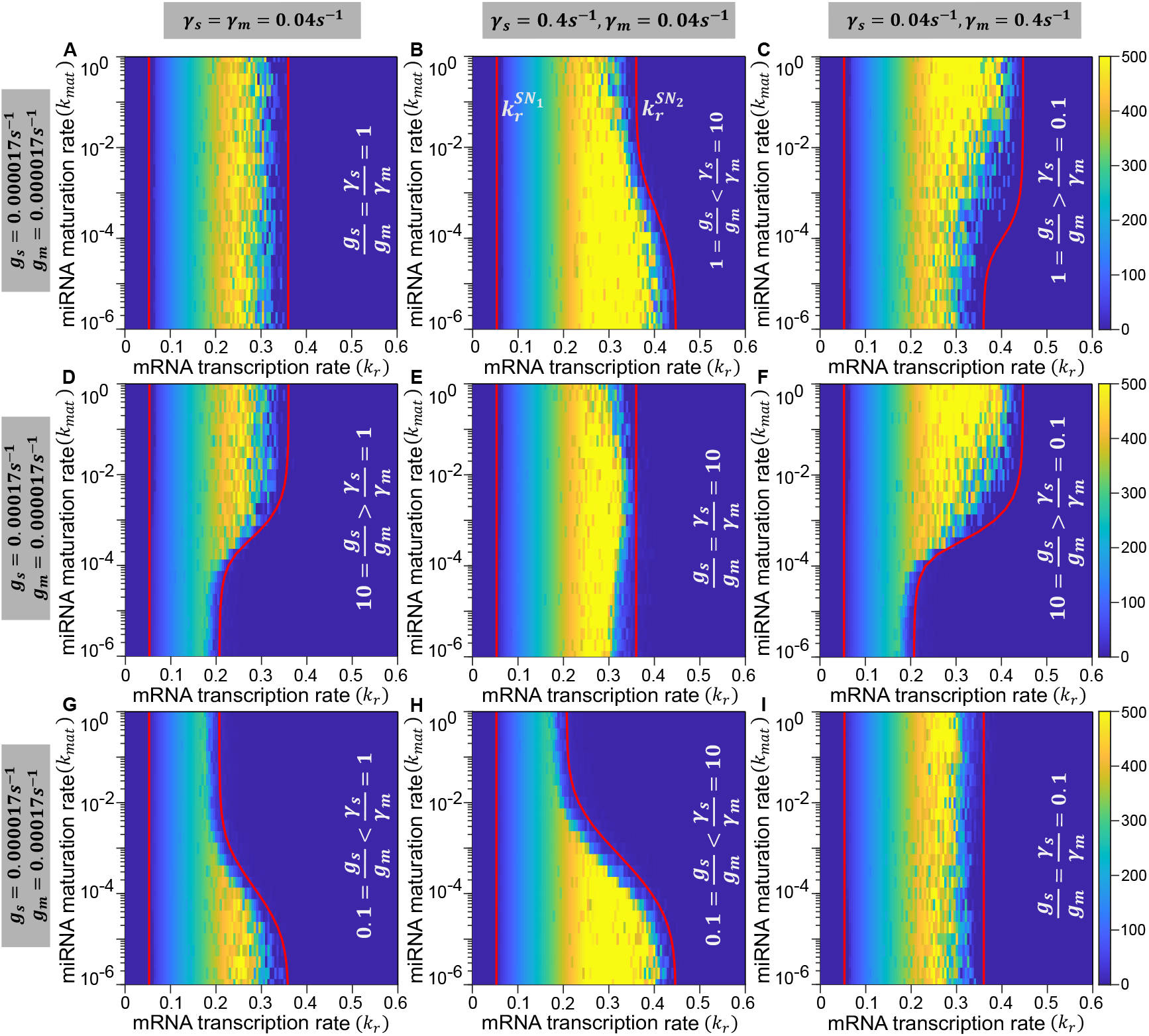
Variation of pre- and mature miRNA degradation rates (*g*_*s*_ and *g*_*m*_) and mRNA-miRNA co-degradation rates (*γ*_*s*_ and *γ*_*m*_) collectively modulates the region of bistability. In each panel, heatmaps show the absolute difference of steady-state mean mRNA, *abs*(⟨*r*⟩) = |⟨*r*⟩_(0,0,0,0)_ − ⟨*r*⟩_(2000,2000,2000,2000)_|, corresponding to zero ((*r*_0_, *p*_0_, *s*_0_, *m*_0_) = (0, 0, 0, 0)) and nonzero ((*r*_0_, *p*_0_, *s*_0_, *m*_0_) = (2000, 2000, 2000, 2000)) initial conditions, obtained from stochastic simulations. Red solid curves in each panel represent the saddle-node bifurcation curves in the *k*_*r*_-*k*_*mat*_ plane, obtained from mean-field stability analysis (see SI Sections 5 and 6). For top row panels (A-C), pre-miRNAs and mature miRNAs have the same degradation rates (*g*_*s*_ = *g*_*m*_). Pre-miRNAs have lower degradation rates (*g*_*s*_ *< g*_*m*_) in middle row panels (D-F), and higher degradation rates (*g*_*s*_ *> g*_*m*_) in the bottom row (G-I), respectively. Note that along the diagonal panels (A, E, I), the realtionship 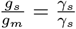 holds and the region of bistability becomes independent of miRNA maturation rate (*k*_*mat*_). When 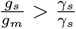, the region of bistability widens as *k*_*mat*_ increases (see panels C, D, F), and when 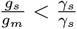, the region of bistability shrinks with *k*_*mat*_ (panels B, G, H). For all panels, *k*_*b*_ = 10^−3^*s*^−1^, *k*_*u*_ = 10^−2^*s*^−1^, 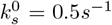, *k*_*s*_ = 0.05*s*^−1^. Other parameters are taken from Table 1.

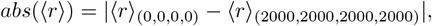

where ⟨*r*⟩_(0,0,0,0)_ and ⟨*r*⟩_(2000,2000,2000,2000)_ are steady-state mean mRNAs obtained from stochastic simulations using two distinct initial conditions: (*r*_0_, *p*_0_, *s*_0_, *m*_0_) = (0, 0, 0, 0) and (*r*_0_, *p*_0_, *s*_0_, *m*_0_) = (2000, 2000, 2000, 2000). Thus, *abs*( ⟨*r*⟩) *>* 0 shows a significant difference in ⟨*r*⟩ for different initial conditions, implying bistability; while *abs*( ⟨*r*⟩) ≈ 0 implies no bistability. The heatmaps of *abs*( ⟨*r*⟩) are plotted in Fig. 3, which shows that the high intensity zones are bounded by the bifurcation boundaries, signifying the agreement between stochastic simulations and mean-field stability analysis.

In Fig. 3A, the individual degradation rates of pre- and mature miRNAs are the same (i.e., *g*_*s*_ = *g*_*m*_), and the miRNA-mRNA co-degradation rates are also the same for both (*γ*_*s*_ = *γ*_*m*_). This implies no essential difference between these two miRNA species. Thus, the region of bistability becomes independent of the pre-miRNA maturation rate, *k*_*mat*_. In Fig. 3B, however, both pre-and mature miRNAs have the same degradation rates, but pre-miRNAs co-degrade with mRNAs faster than mature miRNAs (i.e, *g*_*s*_ = *g*_*m*_ and *γ*_*s*_ *> γ*_*m*_). In this case, when *k*_*mat*_ increases, target mRNAs escape degradation by pre-miRNAs since more pre-miRNAs convert into mature miRNAs. Thus, the mean mRNA, starting from zero, remains at a low level for a shorter range of *k*_*r*_ as *k*_*mat*_ increases, and consequently, the bistable region becomes narrower with *k*_*mat*_ (Fig. 3B). The opposite scenario takes place for *g*_*s*_ = *g*_*m*_ and *γ*_*s*_ *< γ*_*m*_ (Fig. 3C). Here, mature miRNAs codegrade with mRNAs faster than pre-miRNAs, resulting in widening of the bistable region with *k*_*mat*_.

Similarly, Fig. 3D and Fig. 3G show the case of equal co-degradation rates of pre-and mature miRNAs (*γ*_*s*_ = *γ*_*m*_), but unequal individual degradation rates (*g*_*s*_ *> g*_*m*_ or *g*_*s*_ *< g*_*m*_). Here, we reach the same conclusion as previously discussed in Fig. 2. If *g*_*s*_ *> g*_*m*_, pre-miRNAs are more unstable or degradation-prone than mature miRNAs. Thus, increasing *k*_*mat*_ produces more relatively stable mature miRNAs that downregulate mRNAs more effectively, keeping the mean mRNA at a low level for a broader range of *k*_*r*_, and consequently widening the region of bistability (Fig. 3D). The opposite case occurs in Fig. 3G, where the bistable region shrinks with increasing *k*_*mat*_ for *g*_*s*_ *< g*_*m*_.

In general, the relative magnitude of the two ratios, *g*_*s*_*/g*_*m*_ and *γ*_*s*_*/γ*_*m*_, determines how the bistability changes with *k*_*mat*_. Note that *g*_*s*_*/g*_*m*_ encodes the relative lifetime of pre-versus mature miRNAs (when *γ*_*s*_ = *γ*_*m*_); while *γ*_*s*_*/γ*_*m*_ captures relative ability to codegarde target mRNAs by pre- and mature miRNAs (when *g*_*s*_ = *g*_*m*_). Based on the arugemts described above, if 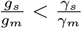, the bistable region shrinks as *k*_*mat*_ increases (as in Fig.3B, G, H). Conversely, when 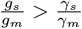, the region of bistability widens with *k*_*mat*_ (see Fig.3 C, D, F).

Interestingly, when *g*_*s*_*/g*_*m*_ = *γ*_*s*_*/γ*_*m*_, the bistable region becomes independent of *k*_*mat*_ (see Fig. 3A, E, I). To understand this, we note that, from the mean-field equations (Eq. 1.a- 1.e), the mean mRNA in the steady-state can be expressed as a function of all rate parameters (see section 7 in SI):

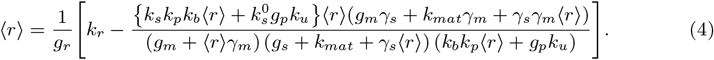

From Eq. 4, considering *g*_*s*_*/g*_*m*_ = *γ*_*s*_*/γ*_*m*_ = *α* we obtain

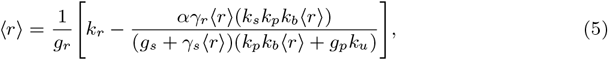

which is independent of *k*_*mat*_. Another intuitive understanding of this *k*_*mat*_ independence follows from the mean-field equations. Adding Eqs. 1.c and 1.d, leads to

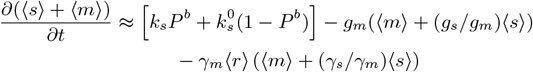

When *g*_*s*_*/g*_*m*_ = *γ*_*s*_*/γ*_*m*_ = *α*, this equation, in the steady-state reduces to

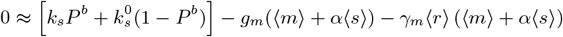

Thus, in the steady-state, 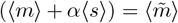 behaves like the mean of a single miRNA species without any *k*_*mat*_ dependence. Overall, *g*_*s*_*/g*_*m*_ and *γ*_*s*_*/γ*_*m*_ act as control parameters in tuning the bistability in the *k*_*r*_-*k*_*mat*_ plane.

#### 3.1.3 mRNA noise peaks and bimodality emerges near bifurcation boundaries

We next study how the miRNA maturation affects the noise in the target mRNAs. Focusing on the parameter regime, 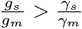 (corresponding to Fig. 3D), we calculate the mRNA Fano factor in the steady-state (defined as variance to mean ratio) from stochastic simulations with two initial conditions: zero and high nonzero initial copy numbers ((*r*_0_, *p*_0_, *s*_0_, *m*_0_) = (0, 0, 0, 0)) and (*r*_0_, *p*_0_, *s*_0_, *m*_0_) = (2000, 2000, 2000, 2000)). The heatmap of the mRNA Fano factor for zero initial condition (Fig. 4A) shows that high intrinsic noise emerges near the right saddle-node bifurcation boundary 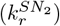, while the mRNA noise peaks around the left bifurcation boundary 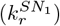 corresponding to the nonzero initial condition (Fig. 4C). Accordingly, Fano factors at a fixed maturation rate *k*_*mat*_ have maxima near the bifurcation boundaries (Fig. 4B, 4D). For both initial conditions, we also observe that peak positions in the Fano factor shift with the maturation rate, *k*_*mat*_, and the peak heights decrease with increasing *k*_*mat*_, showing that the noise level decreases with *k*_*mat*_ (Fig. 4B, 4D). The opposite scenario happens for 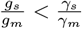, where mRNA noise level increases with *k*_*mat*_ (see Fig. S1).

**Figure 4.**
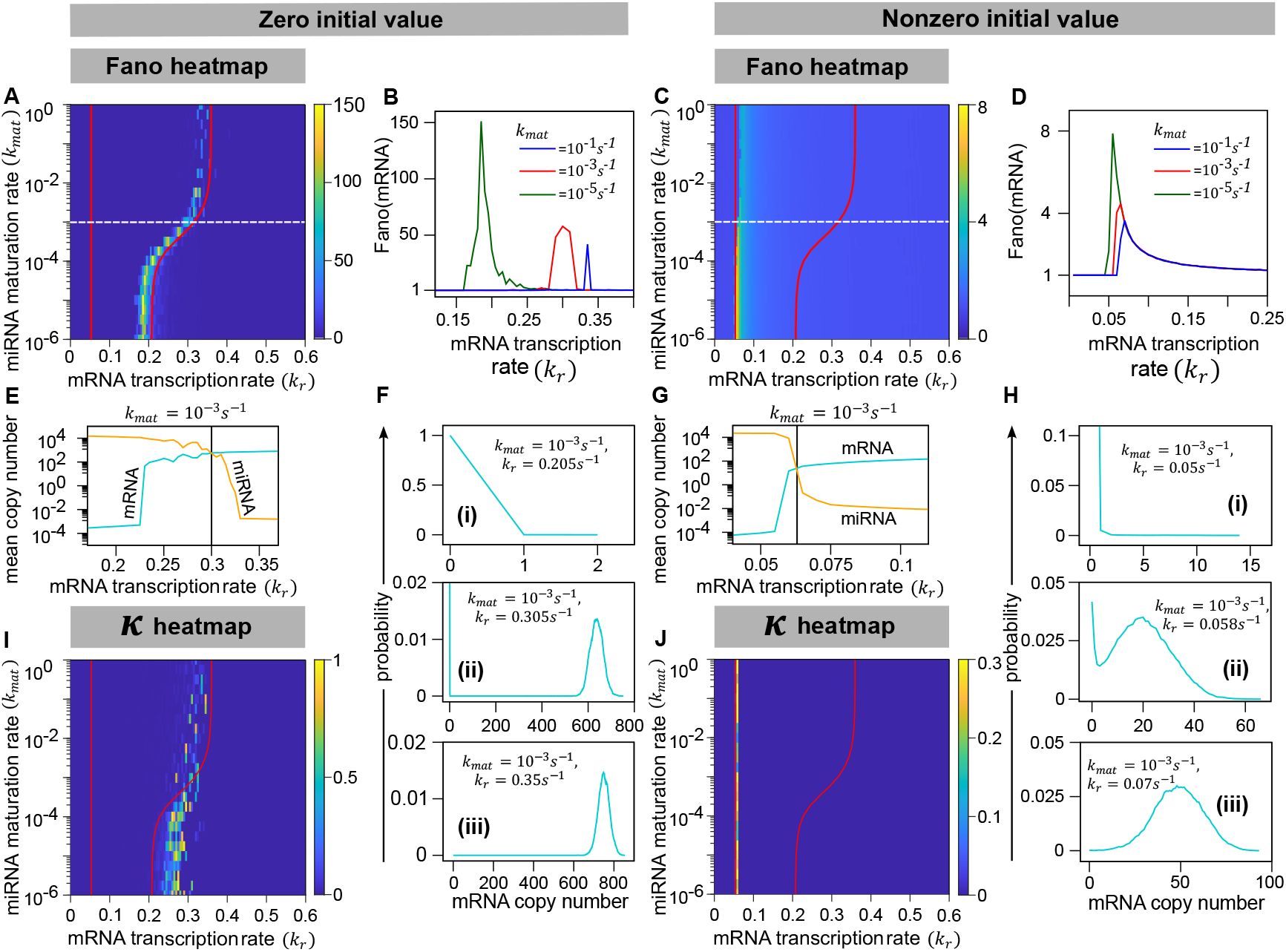
Initial condition-dependent noise amplification and bimodality near bifurcation boundaries in miRNA-mediated positive feedback. Results are shown from stochastic simulations with two distinct initial conditions: Zero (panels A, B, E, F, I) and high nonzero (C, D, G, H, J) initial copy numbers, i.e., (*r*_0_, *p*_0_, *s*_0_, *m*_0_) = (0, 0, 0, 0) and (*r*_0_, *p*_0_, *s*_0_, *m*_0_) = (2000, 2000, 2000, 2000), respectively. (A, C) Heatmaps of the steady-state Fano factor of mRNAs are shown in the *k*_*r*_-*k*_*mat*_ plane, for zero (A) and nonzero (C) initial conditions. Red solid curves represent the saddle-node bifurcation boundaries obtained from stability analysis of mean-field equations (see SI, sections 5-6). (B, D) The steady-state Fano factors of mRNA, corresponding to different maturation rates (*k*_*mat*_), are plotted against *k*_*r*_. (E, G) At a fixed *k*_*mat*_, the mean mRNA ( ⟨*r*⟩) and total mean miRNA ( ⟨*s*⟩ + ⟨*m*⟩) show transitions as functions of *k*_*r*_, for both zero (E) and nonzero (G) initial conditions. Dashed horizontal lines in A and C represent *k*_*mat*_ = 10^−3^*s*^−1^, for which means are plotted in E and G. Black vertical lines in E and G represent the positions of bifurcation points. (F, H) For *k*_*mat*_ = 10^−3^*s*^−1^, steady-state mRNA distributions spanning the right (F(i-iii)) and left (H(i-iii)) bifurcation boundaries are shown. Note the bimodal distributions near the right and left bifurcation boundaries for zero and nonzero initial conditions, respectively (F(ii) and H(ii)). (I, J) The heatmaps of bimodality strength (*κ*) are shown in the *k*_*r*_ vs *k*_*mat*_ plane. The same red bifurcation curves, as in A and C, are replotted in I and J. For all panels, *k*_*b*_ = 10^−3^*s*^−1^, *k*_*u*_ = 10^−2^*s*^−1^, 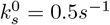, *k*_*s*_ = 0.05*s*^−1^, *g*_*s*_ = 0.00017*s*^−1^, *g*_*m*_ = 0.000017*s*^−1^, *γ*_*s*_ = *γ*_*m*_ = 0.04*s*^−1^. Other parameters are from Table 1.

To understand the emergence of noise amplification near the bifurcation boundaries, we note that the mean mRNA ( ⟨*r*⟩) and mean total miRNAs ( ⟨*s*⟩ + ⟨*m*⟩) cross over each other near the bifurcation boundary (Fig. 4E, 4G). For both initial conditions, away from the bifurcation boundary, either mRNAs or miRNAs (i.e., the sum of pre- and mature miRNAs) dominate over each other on average; in contrast, near the bifurcation boundary, mRNAs and miRNAs are comparable to each other, and they are present in low copy numbers (Fig. 4E, 4G). Thus, mRNAs are either stochastically codegraded by miRNAs or escape degradation and remain free, generating high fluctuations in their copy numbers near the bifurcation point. Correspondingly, the steady-state mRNA distribution becomes bimodal near the bifurcation point (Figs. 4F(ii) and 4H(ii)), suggesting that substantial noise can create two distinct subpopulations: mRNAs can be present in either low or high copy numbers across many genetically identical cells. However, the mRNA distribution becomes bell-shaped above the bifurcation point, and it becomes exponential-like below the bifurcation point (Figs. 4F, 4H).

To further investigate emergence of bimodal mRNA distributions across the bifurcation boundary, we calculate the ‘bimodality strength’ of mRNA distributions [45], defined as *κ* = (*H*_*low*_ − *H*_*valey*_)*/H*_*high*_, where *H*_*low*_, *H*_*valley*_ and *H*_*high*_ denote the heights of the lower mode, valley region, and higher mode, respectively (note, 0 ≤ *κ* ≤ 1). Thus, *κ* = 0 means unimodality, while *κ >* 0 means bimodality. Figs. 4I and 4J show the heatmaps of *κ* in the *k*_*r*_ vs *k*_*mat*_ plane. For both initial conditions, the high-intensity regions in the heatmaps of mRNA Fano factor and *κ* overlap around the saddle-node bifurcation boundaries (Figs. 4A, 4I; and Figs. 4C, 4J for zero and nonzero initial conditions, respectively). Thus, the mRNA noise, sensitive to the initial condition, can induce bimodality near bifurcation boundaries. Note that high-intensity regions in the Fano factor and *κ* heatmaps only appear around the bifurcation boundaries, not within the whole bistability region. Thus, contrary to the naïve expectation, deterministic bistability may not always lead to noise-induced bimodality. This raises a question: Does the ‘deterministic bistability’ and noise-induced ‘stochastic bimodality’ always appear together in the overlapping region of the parameter space?

#### 3.1.4 Feedback strength modulates deterministic bistability and stochastic bimodality, which may not co-occur

To investigate whether ‘deterministic bistability’ (DB) and ‘stochastic bimodality’ (SB) always appear together, we vary the positive feedback strength, defined as the ratio of rates of protein binding/unbinding with the miRNA gene (*β* = *k*_*b*_*/k*_*u*_). We focus on the case, 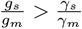 and discuss here the simulation results with the zero initial condition, i.e., (*r*_0_, *p*_0_, *s*_0_, *m*_0_,) = (0, 0, 0, 0) (corresponding to Fig. 4A).

The heatmaps of bimodality strength (*κ*) obtained from stochastic simulations are shown in Fig. 5A-5E, along with the bifurcation boundaries (denoted by 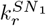 and 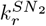) obtained from deterministic stability analysis. The DB region is bounded by the bifurcation boundaries. We find that, for intermediate feedback strengths (*β* = 10^−5^, 10^−1^, and 1), both DB and SB appear together near the right bifurcation boundary (Fig. 5B-5D). The regions of DB and SB overlap but are not exactly the same. We also observe that both DB and SB regions shift from high to low mRNA transcription rates (*k*_*r*_) as the feedback strength (*β*) increases (Fig. 5B-5D). Interestingly, DB disappears for the limiting cases of very low feedback (*β* = 10^−7^) and extremely high feedback strengths (*β* = 10^3^), although the SB persists (Fig. 5A and E). In both limiting cases, the mean mRNA still exhibits a transition from low to high values as *k*_*r*_ increases, and the mRNA noise also peaks near the transition region (see Fig. S2); however, there is no bistability predicted from deterministic analysis. Thus, in the limiting cases, bimodality emerges solely due to stochastic fluctuations, rather than any underlying bistability.

**Figure 5.**
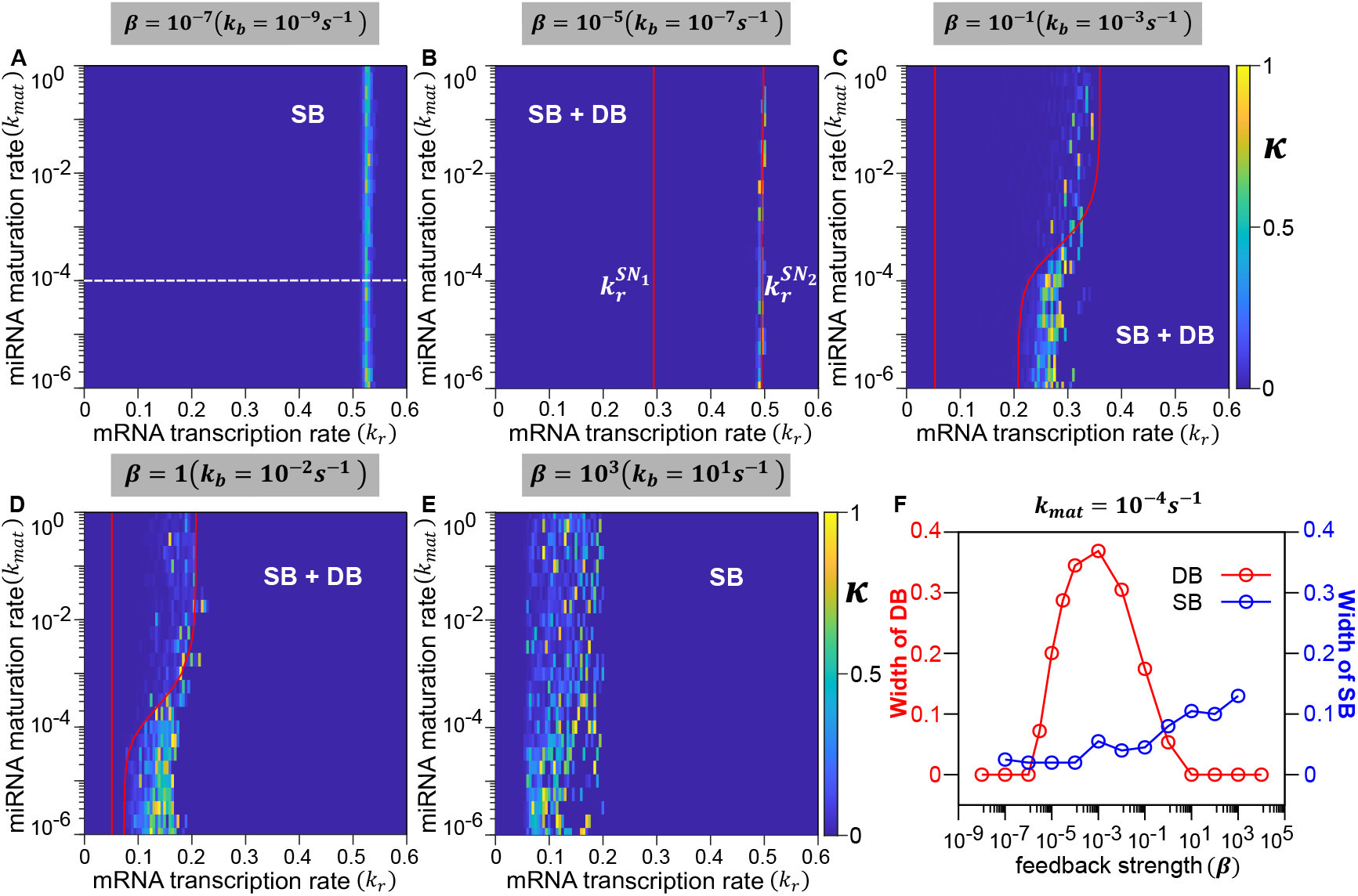
Effect of varying feedback strength on deterministic bistability (DB) and stochastic bimodality (SB) in a positive feedback loop. (A-E) The heatmaps of bimodality strength (*κ*) obtained from stochastic simulations with the zero initial condition ((*r*_0_, *p*_0_, *s*_0_, *m*_0_) = (0, 0, 0, 0)) are shown in the *k*_*r*_-*k*_*mat*_ plane. Red solid curves in B, C, and D are saddle-node bifurcation boundaries obtained from mean-field analysis (denoted by 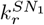 and 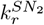 in B). Note that the DB (i.e., bifurcation curves) does not exist in A and E. (F) The widths of DB and SB regions are plotted against the feedback strength (*β*) for a fixed pre-miRNA maturation rate. The dashed horizontal line in A represents the fixed maturation rate, *k*_*mat*_ = 10^−4^*s*^−1^, along which the widths of DB and SB are plotted in F. For all panels, *k*_*u*_ = 10^−2^*s*^−1^, 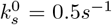, *k*_*s*_ = 0.05*s*^−1^, *g*_*s*_ = 0.00017*s*^−1^, *g*_*m*_ = 0.000017*s*^−1^, *γ*_*s*_ = *γ*_*m*_ = 0.04*s*^−1^. Other parameters are taken from Table 1.

To compare the regions of DB and SB, we calculate the ‘width of bistability’ defined as the difference between the right and left bifurcation points at a fixed miRNA maturation rate, *k*_*mat*_ (given by 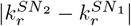). Similarly, we define the ‘width of stochastic bimodality’ as the length of a region with nonzero *κ* values at a fixed *k*_*mat*_, i.e., the width of SB is given by 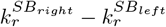, where 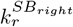 and 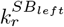 are points on the *k*_*r*_-axis (at a fixed *k*_*mat*_), within which *κ* is nonzero in the *κ*-heatmaps. Fig. 5F shows both the widths of DB and SB for a fixed miRNA maturation rate (*k*_*mat*_ = 10^−4^*s*^−1^) as functions of the feedback strength, *β*. We find that the width of the DB exhibits non-monotonic behavior with *β*, indicating that the DB region does not exist at extremely low and high feedback strengths, but widens for intermediate feedback strengths. In contrast, the width of the SB region increases monotonically with increasing feedback strength. Therefore, the feedback strength modulates both DB and SB in the target gene expression. However, DB does not always coincide with SB that arises due to noise amplification near the transition region from low to high expression level.

### 3.2 Effects of miRNA Maturation on the Target Gene Expression in a Negative Feedback Loop

#### 3.2.1 Negative feedback strength and miRNA maturation tune the oscillatory expression

Some theoretical studies of miRNA-dependent negative feedback, although not incorporating miRNA maturation, have shown that negative feedback can lead to oscillations in target gene expression [56, 120]. To investigate how miRNA maturation affects the oscillatory dynamics, we choose the parameter regime, *g*_*s*_ = *g*_*m*_ and *γ*_*s*_ *< γ*_*m*_ (i.e., 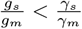). We have numerically solved the mean-field equations (Eq. 1.a-1.e) in the steady-state and plotted the mean mRNA (⟨*r*⟩) in Fig. 6A-C as a function of mRNA transcription rate (*k*_*r*_) and miRNA maturation rate (*k*_*mat*_), for different negative feedback strengths. As shown in previous models without miRNA maturation [1, 73, 76], the mean mRNA shows a transition from low to high expression beyond a threshold value of *k*_*r*_. Interestingly, oscillation only emerges for intermediate feedback strength (Fig. 6B, where the orange region represents the oscillatory regime). Moreover, oscillation disappears with increasing miRNA maturation rate (Fig. 6B).

**Figure 6.**
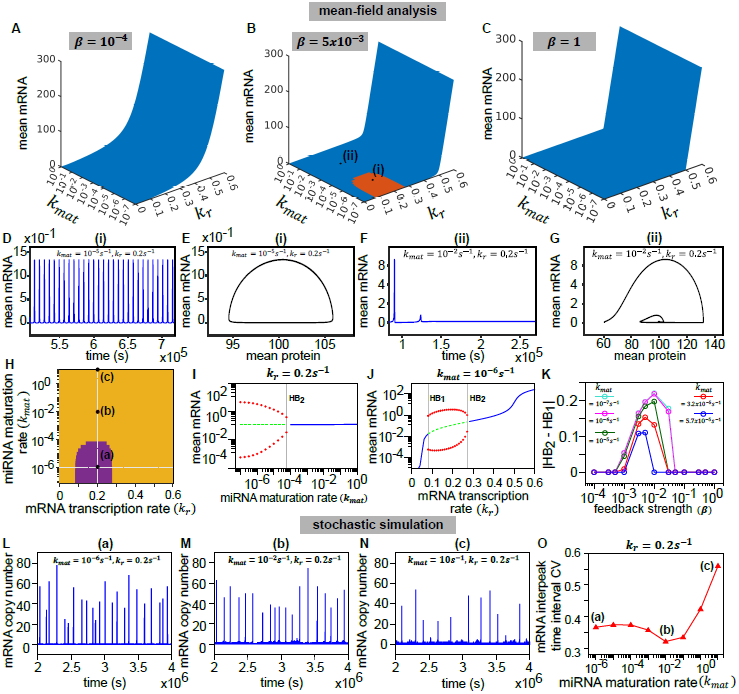
Feedback strength and pre-miRNA maturation tune the oscillatory gene expression in a miRNA-mediated negative feedback loop. (A-C) The mean mRNAs (obtained from mean-field solutions in the steady state) are shown in the *k*_*r*_ vs *k*_*mat*_ plane for different feedback strengths. The orange region in B and the blue regions in A-C indicate oscillatory and non-oscillatory regimes, respectively. (D-G) The steady-state time profiles of mean mRNAs (D, F) and phase portraits in the plane of mean mRNA vs mean protein (E, G) are shown for points (i) and (ii), corresponding to Panel B. Sustained oscillations and limit cycles are seen for point (i) (D, E), but not for point (ii) (F, G). (H) The non-oscillatory (yellow) and oscillatory (purple) regions are replotted from Panel B in the *k*_*r*_-*k*_*mat*_ plane. Vertical and horizontal dashed lines indicate the variation of *k*_*mat*_ or *k*_*r*_, along which the bifurcation diagrams are shown in I and J. (I-J) Steady-state mean mRNAs with varying *k*_*mat*_ (I) and varying *k*_*r*_ (J), where *HB*_1_ and *HB*_2_ denote Hopf bifurcation points. In I and J, blue solid curves represent the ‘stable’ solutions and red circles represent the maxima and minima of mean mRNAs, surrounding an ‘unstable’ state (denoted by green dashed curves). For stability analysis, see Section 6 of SI. (K) The ‘width of oscillatory region’ ( |*HB*_2_ − *HB*_1_|) is plotted with varying feedback strengths. (L-N) The steady-state stochastic trajectories of mRNAs obtained from simulations. (O) The CV of mRNA interpeak time interval (*CV* (*τ*_*peak*_)), obtained from stochastic oscillations, is plotted as a function of miRNA maturation rate (*k*_*mat*_). The (*k*_*r*_, *k*_*mat*_) coordinates of the points **(a), (b)**, and **(c)** in H are, **(a)** *k*_*r*_ = 0.2*s*^−1^, *k*_*mat*_ = 10^−6^*s*^−1^, **(b)** *k*_*r*_ = 0.2*s*^−1^, *k*_*mat*_ = 10^−2^*s*^−1^, and **(c)** *k*_*r*_ = 0.2*s*^−1^, *k*_*mat*_ = 10*s*^−1^, respectively. Parameters: *k*_*u*_ = 10^−2^*s*^−1^, 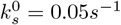, *k*_*s*_ = 0.5*s*^−1^, *g*_*s*_ = *g*_*m*_ = 0.000017*s*^−1^, *γ*_*s*_ = 4*s*^−1^, *γ*_*m*_ = 0.004*s*^−1^. For panels (D-J) and (L-O), *k*_*b*_ = 5x10^−5^*s*^−1^(*β* = 5x10^−3^). Other parameters are from Table 1.

To understand the origin of oscillations, note that for intermediate feedback strengths (quantified by the protein binding–unbinding ratio, *β* = *k*_*b*_*/k*_*u*_), the miRNA gene toggles between protein-bound and unbound states. However, for very high or low *β*, it remains mostly in one state. When the miRNA maturation rate (*k*_*mat*_) is low, free pre-miRNAs accumulate, exceeding the abundance of mature miRNAs. In this regime, if pre-miRNAs co-degrade with target mRNAs faster than mature miRNAs (*γ*_*s*_ *> γ*_*m*_), they strongly reduce mRNA levels, leading to reduced protein production. This drop in protein switches the miRNA gene to the unbound state, where miRNA synthesis is lower (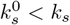 for negative feedback). Reduced miRNA synthesis then allows mRNAs to rise again, increasing protein levels and driving the miRNA gene back to the protein-bound state, where miRNA synthesis is higher. This repeating cycle of lower and higher miRNA synthesis creates sustained oscillations in mean mRNA levels.

However, this alternating cycle in miRNA synthesis breaks down when the miRNA maturation rate (*k*_*mat*_) is high. In this regime, even if mRNAs are more prone to be co-degraded by pre-miRNAs than by mature miRNAs (*γ*_*s*_ *> γ*_*m*_), the pool of free pre-miRNAs becomes too small to strongly downregulate target mRNAs. As a result, the protein level stays high primarily, and the miRNA gene hardly switches between bound and unbound states. Thus, the oscillations in mean mRNA levels disappear. In the opposite regime, when *g*_*s*_ = *g*_*m*_ and *γ*_*s*_ *< γ*_*m*_ (i.e., when 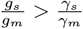) oscillations emerge for intermediate feedback strengths if the maturation rate is high (Fig. S3).

Inside the oscillatory regime (denoted by (i) in the *k*_*r*_-*k*_*mat*_ space in Fig. 6B), we observe sustained oscillation in the steady-state time profile of mean mRNA (Fig. 6D), and correspondungly, a limit cycle in the phase space of the mean mRNA (⟨*r*⟩) versus mean protein (⟨*p*⟩) (Fig. 6E). Away from the oscillatory regime (for point (ii) in Fig. 6B), the time profile of ⟨*r*⟩ shows damped oscillations (Fig. 6F), producing a spiral in the ⟨*r*⟩-⟨*p*⟩ phase space. (Fig. 6G). Similar transitions from sustained to damped oscillations can be seen when the transcription rate is varied at a fixed *k*_*mat*_ (Fig. S4).

The boundary of the oscillatory regime (purple region in Fig. 6H, reproduced from 6B) can be determined by numerically finding the positions of Hopf bifurcations from the maxima and minima of the mean-field solution of ⟨*r*⟩. For a fixed *k*_*r*_ (along the vertical dashed line in Fig. 6H), ⟨*r*⟩ exhibits a ‘sub-critical’ Hopf bifurcation (Fig. 6I) as miRNA maturation rate (*k*_*mat*_) increases. Similarly, for a fixed *k*_*mat*_ (along the horizontal dashed line in Fig. 6H), ⟨*r*⟩ exhibits ‘super-critical’ (denoted by *HB*_1_) and ‘sub-critical’ (denoted by *HB*_2_) Hopf bifurcations with varing *k*_*r*_ (Fig. 6J). To quantify the effect of miRNA maturation on the oscillatory regime, we define the quantity |*HB*_2_ − *HB*_1_| (i.e., the difference between the positions of ‘super-critical’ and ‘sub-critical’ Hopf bifurcation points) as the ‘width of oscillatory region’ (note, |*HB*_2_ − *HB*_1_| ∼ 0 implies no oscillation). In Fig. 6K, we show |*HB*_2_ − *HB*_1_| for different miRNA maturation rates, as a function of the negative feedback strength (*β*). We observe non-monotonic behavior of |*HB*_2_ − *HB*_1_| versus *β* curves, reflecting the emergence of oscillatory dynamics only for an intermediate range of *β*. Additionally, the width of the oscillatory regime decreases with increasing *k*_*mat*_ (Fig. 6K), indicating that the negative feedback strength and miRNA maturation rate jointly modulate oscillation in target gene expression.

We next compare our mean-field prediction of oscillatory dynamics with the exact stochastic simulations. A natural question is whether noise can disrupt oscillations predicted by the mean-field analysis. To investigate this, we performed stochastic simulations for different miRNA maturation rates (for points (a), (b), and (c) along the dashed vertical line in Fig. 6H). The trajectories of mRNAs show stochastic oscillations inside the oscillatory regime, as expected (Fig. 6L). However, note that the stochastic oscillation visibly remains regular even outside the mean-field oscillatory regime (see Fig. 6M corresponding to point (b) in Fig. 6H). But it ultimately becomes irregular and noisy very far away from the oscillatory regime (Fig. 6N).

To probe the regularity in the noisy oscillations, we quantify the noise in the time period, given by the coefficient of variation (CV) of the mRNA interpeak time interval (*τ*_*peak*_) in the steady-state: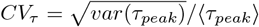 . A low *CV*_*τ*_ implies regularity in the appearance of peaks in the mRNA trajectories. Fig. 6O shows the calculated *CV*_*τ*_ as a function of the miRNA maturation rates, *k*_*mat*_ (along the vertical dashed line in Fig. 6H). Notably, the values of *CV*_*τ*_ remain low even outside the deterministic oscillatory regime, suggesting that regularity in stochastic oscillations persists beyond the mean-field prediction (see point (b) in Fig. 6O). Thus, the stochastic oscillatory regime does not necessarily overlap with the deterministic limit cycle regime. We come to this same conclusion by measuring *CV*_*τ*_ for varying transcription rate at a fixed *k*_*mat*_ (Fig. S5).

#### 3.2.2 Noise-induced stochastic bimodality emerges near the expression threshold outside the oscillatory regime

We next examine how miRNA-mediated negative feedback shapes mRNA distributions. For an intermediate feedback strength (parameters as in Fig. 6B), we computed the mean mRNA level, ⟨*r*⟩, and the mRNA Fano factor (*var*(*r*)*/* ⟨*r*⟩) across the *k*_*r*_-*k*_*mat*_ parameter space (Fig. 7A, B). Consistent with mean-field predictions (Fig. 6B), ⟨*r*⟩ exhibits a sharp threshold separating low and high expression regimes (Fig. 7A). This transition region is accompanied by enhanced mRNA noise (Fig. 7B). To assess whether this enhanced noise leads to bimodality, we quantified the bimodality strength, *κ*, of the mRNA distributions (Fig. 7C). High *κ* values are localized near the threshold (see point (b) in Fig. 7C), indicating noise-induced bimodality.

**Figure 7.**
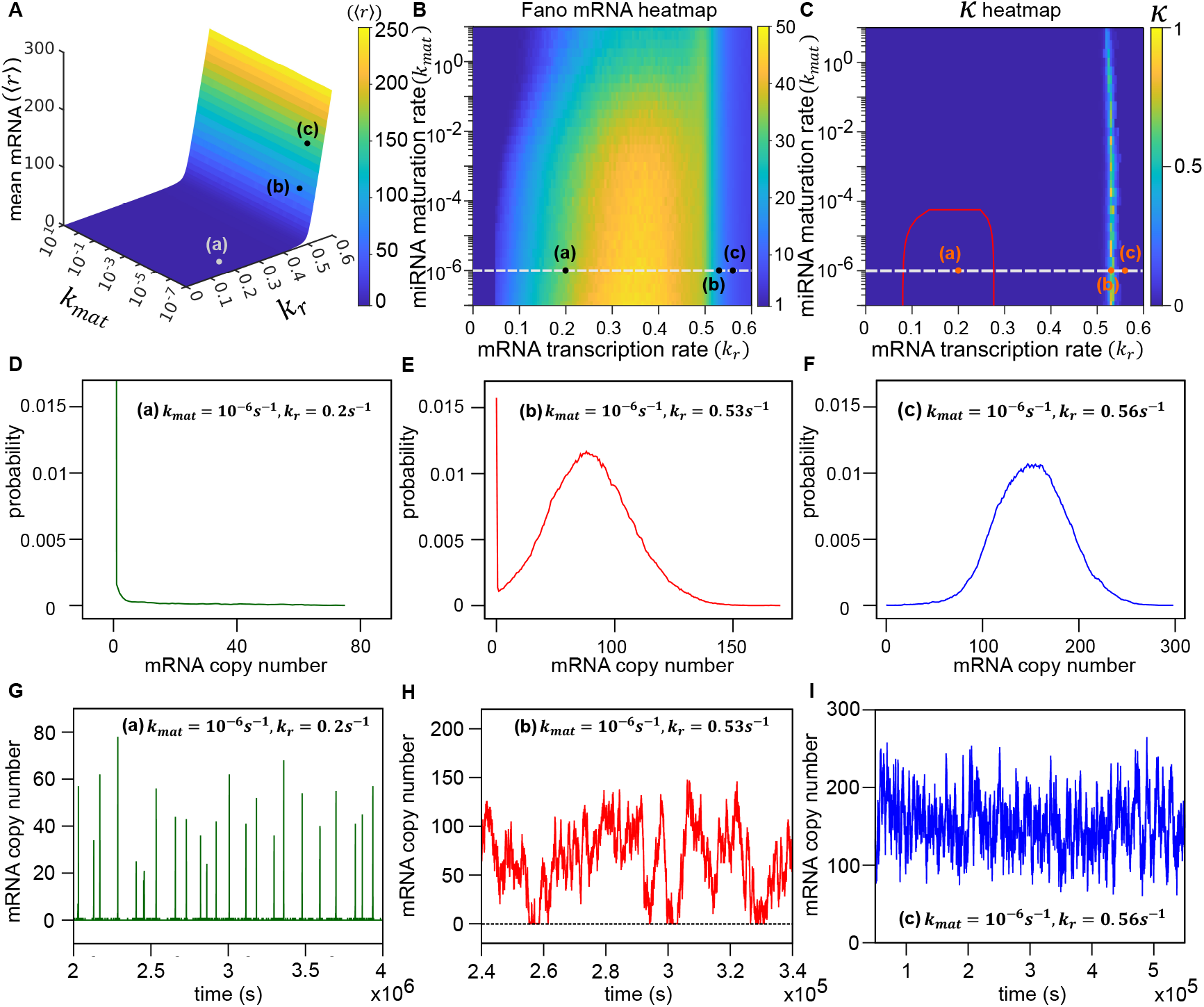
In a negative feedback loop, bimodal mRNA distributions emerge near the mRNA threshold outside the oscillatory region. (A-C) Heatmaps of mean mRNA, ⟨*r*⟩ (A), Fano factor of mRNA (B), and bimodality strength, *κ* (C), obtained from stochastic simulations for an intermediate feedback strength (*β* = 5x10^−3^). In B and C, the horizontal dashed line represents *k*_*mat*_ = 10^−6^*s*^−1^. In C, the area enclosed by the solid red curve represents the oscillatory region (reproduced from Fig. 6H) obtained from mean-field analysis. The (*k*_*r*_, *k*_*mat*_) coordinates of the three highlighted points in A-C are: (a) *k*_*r*_ = 0.2*s*^−1^, *k*_*mat*_ = 10^−6^*s*^−1^, (b) *k*_*r*_ = 0.53*s*^−1^, *k*_*mat*_ = 10^−6^*s*^−1^, and (c) *k*_*r*_ = 0.56*s*^−1^, *k*_*mat*_ = 10^−6^*s*^−1^ (D-F) Steady-state mRNA distributions, obtained for the points (a), (b), and (c). (G-I) Stochastic trajectories of mRNA copy numbers, obtained from simulations, are shown for the points (a), (b), and (c). For all panels, *k*_*b*_ = 5x10^−5^*s*^−1^, *k*_*u*_ = 10^−2^*s*^−1^(*β* = 5x10^−3^), 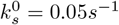, *k*_*s*_ = 0.5*s*^−1^, *g*_*s*_ = *g*_*m*_ = 0.000017*s*^−1^, *γ*_*s*_ = 4*s*^−1^, *γ*_*m*_ = 0.004*s*^−1^. Other parameter values are taken from Table 1.

To elucidate the origin of this bimodality and if it is connected to oscillatory dynamics, we analyzed mRNA distributions at three representative points in the paramter space (along the dashed line in Fig. 7C): inside the deterministic limit-cycle regime (a), near the threshold where *κ* is maximal (b), and away from the threshold (c). The steady-state mRNA distributions evolve from an exponential-like form (a; Fig. 7D) to a bimodal distribution near the threshold (b; Fig. 7E), and finally to a unimodal, bell-shaped distribution away from the threshold (c; Fig. 7F). Stochastic time trajectories reveal that, near the threshold, mRNA copy numbers stochastically switch between near-zero and high values in a bursty manner (Fig. 7H), generating bimodality. In contrast, away from the threshold, mRNA levels remain persistently low or high (Fig. 7G, 7I). As previously shown [1], this behavior arises from stochastic toggling of the miRNA gene between protein-bound and unbound states near the threshold, leading to two distinct miRNA synthesis rates. Away from the threshold, this switching is suppressed, and miRNAs are produced mainly at a higher or a lower rate. Thus, stochastic bimodality can emerge purely from noise amplification near the threshold, outside the deterministic oscillatory regime.

## 4 Discussion

Our study develops a stochastic gene regulatory model that explicitly incorporates miRNA maturation and the resulting competition between precursor miRNAs (pre-miRNAs) and mature miRNAs for a shared target mRNA pool. By extending earlier miRNA-dependent feedback models [120] to include pre-miRNA dynamics, we systematically examine how the maturation rate, relative degradation rates of mature versus pre-miRNAs, and miRNA–mRNA co-degradation rates reshape both deterministic behavior (such as bistability and oscillations) and stochastic outcomes (noise amplification and bimodal mRNA distributions).

At the mean-field level, our model shows that miRNA-mediated positive feedback generates bistability via saddle-node bifurcations, while negative feedback produces oscillations through Hopf bifurcations, consistent with previous theoretical studies that neglected maturation dynamics [1, 9, 32, 56, 76]. The key advancement is the demonstration that miRNA maturation and competition between mature and pre-miRNAs quantitatively modulate the extent of these dynamical regimes, sometimes expanding or shrinking bistable and oscillatory regions in parameter space (Fig. 2, Fig. 3, and Fig. 6B). This identifies maturation as a crucial control knob for feedback-driven dynamics, rather than a kinetic detail.

Beyond deterministic predictions, stochastic simulations reveal that bimodal mRNA distributions emerge near bifurcation boundaries in the positive feedback, but also appear outside the regions of deterministic bistability (Fig. 5). Thus, stochastic bimodality does not always occur with deterministic bistability. Similarly, for the negative feedback, stochastic bimodality in mRNA distributions arises near expression thresholds, outside regions of limit cycles (Fig. 7C). While noise-induced bimodality in miRNA-mediated networks has been reported previously [1, 22, 32, 76], our work demonstrates that competition between precursor and mature miRNAs substantially modulates the parameter space supporting bimodality. In both positive and negative feedback loops, stochastic bimodality coincides with enhanced intrinsic noise (Fig. 4, Fig. 7) but does not necessarily overlap with mean-field bifurcation boundaries. These results reinforce the earlier concept that stochastic bimodality cannot be inferred solely from deterministic analysis [32, 114], while extending this to another biologically detailed regulatory circuit.

Although prior models have predicted noise-induced bimodality in miRNA-mediated feedback [22, 32, 76], our present work differs by explicitly modeling pre-miRNA species and maturation-dependent competition, motivated by experimental evidence that precursor miRNAs can bind targets and regulate expression [84, 99, 113]. This extra layer of regulation enables us to make new, quantitative predictions regarding how maturation kinetics reshape bistability, oscillatory, and bimodal behavior, which cannot be obtained from models that assume a single effective miRNA pool.

Intriguingly, experiments by Bosia et al. [10] have demonstrated that competition between two target mRNAs sharing the same miRNA pool results in bimodal distributions of target expression. Their experimental system uses bidirectional plasmids cotransfected into mammalian cells, where the plasmid expresses a fluorescent reporter gene along with engineered miRNA target sites. The fluorescent outputs served as readouts for expression levels and were quantified by flow cytometry to measure the expression distribution. Such a setup for controlled variation of target expression may be utilized to check our model predictions. For instance, genetic up- or downregulation of Dicer and associated proteins that process pre-miRNAs into mature miRNAs can demonstrate the effect of miRNA maturation on target expression distribution, allowing the verification of our model.

Overall, our study provides a mechanistically grounded extension of existing miRNA feedback models, clarifying how miRNA maturation and precursor competition influence stochastic gene expression. By comparing deterministic dynamical regimes with target mRNA distributions, it offers testable predictions on when bimodal expression could arise even in the absence of bistability or oscillations at the mean-field level. Our analysis of single-target miRNA feedback motifs paves the way for extending the framework to multi-target or network-level regulation, which remains an open future direction.

## Materials and methods

We simulated the model (Fig.1) using Gillespie algorithm [33]. The codes were written in FORTRAN90, and data analysis was done using MATLAB. Codes are available at the following link: https://github.com/PhyBi/miRNA-maturation-project

### Parameter choices

#### mRNA and miRNA transcription rates (*k*_*r*_, 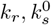 and *k*_*s*_)

Following Bosia et al. [9], Noorbakhsh et al. [76], and Adhikary et al. [1], we varied the transcription rates of mRNA from 0*s*^−1^ to 0.6*s*^−1^ and miRNA transcription rates were taken as 0.5*s*^−1^ and 0.05*s*^−1^ for elevated and basal states (see Table 1).

#### Protein translation rate (*k*_*p*_)

In Wang et al. [120] protein translation rate (*k*_*p*_) is taken to be 1*min*^−1^ ≈ 0.017*s*^−1^, which was assumed in table 1.

#### mRNA and protein degradation rates (*g*_*r*_ and *g*_*p*_)

The protein half-lives are usually longer than those of their mRNA transcripts for different eukaryotic cells (see Jia et al. [44, 46]). In general, mRNA degradation rates are an order of magnitude higher than protein degradation rates in eukaryotes. The protein degradation rate of budding yeast, fission yeast, and human cells is similar [44]: *g*_*p*_ ∼ 0.00002*s*^−1^. The mRNA degradation rates in budding yeast, fission yeast, and human cells are 0.0002*s*^−1^, 0.0004*s*^−1^, and 0.0006*s*^−1^, respectively. Thus, we considered the average value *g*_*r*_ = 0.0004*s*^−1^, i.e., *g*_*r*_ ∼ 10*g*_*p*_.

#### Protein binding and unbinding rate of miRNA-coding gene (*k*_*b*_ and *k*_*u*_)

We assumed the protein binding and unbinding rates from Adhikary et al. [1].

#### Degradation rates of pre-miRNA and mature miRNAs (*g*_*s*_ and *g*_*m*_)

The median half-life of mature miRNAs (for the strands that bind to Ago proteins) is approximately 11.4*h* [83, 122]. Therefore, we consider the degradation rate of mature miRNA to be *g*_*m*_ ∼ *ln*2*/*11.4*h*^−1^ ≈ 0.000017*s*^−1^. We also assume that the degradation rates of pre-miRNAs are similar to those of mature miRNAs, i.e., *g*_*s*_ ∼ *g*_*m*_. Nevertheless, we varied the degradation rates of pre- and mature miRNAs (*g*_*s*_ and *g*_*m*_) over orders of magnitude up to a similar order of *g*_*r*_.

#### Co-degradation rates of target mRNA with pre- and mature miRNAs (*γ*_*s*_ and *γ*_*m*_)

As there is no experimental measurement (to the best of our knowledge), we varied both the rates over orders of magnitude (see Table 1).

## Supporting information

Impact of competition between precursor and mature microRNAs on stochastic gene expression

## Acknowledgment

RA thanks the CSIR, Government of India, for his fellowship (award no. 09/921(0282)/2019-EMR-I). DD acknowledges the financial support from IISER Kolkata and ANRF, Govt. of India (Grant No. ANRF/ARGM/2025/001984/TS and EEQ/2023/000551). We also thank Dr. MK Jolly (IISC Bangalore, India) for useful discussions.

## AUTHOR DECLARATIONS

### Author Contributions

Simulations, calculations, and data analysis were performed by RA. The study was supervised by DD, and the paper was written jointly by RA and DD.

## DATA AVAILABILITY

Simulation codes are available at the link: https://github.com/PhyBi/miRNA-maturation-project

