## Supplementary material for "Impact of competition between precursor and mature microRNAs on stochastic gene expression": Impact of competition between precursor and mature microRNAs on stochastic gene expression

January 3, 2026

**1** Department of Biological Sciences, Indian Institute of Science Education And Research Kolkata, Mohanpur, Nadia 741246, West Bengal, India.

†
‡

#### Supplemental Figures

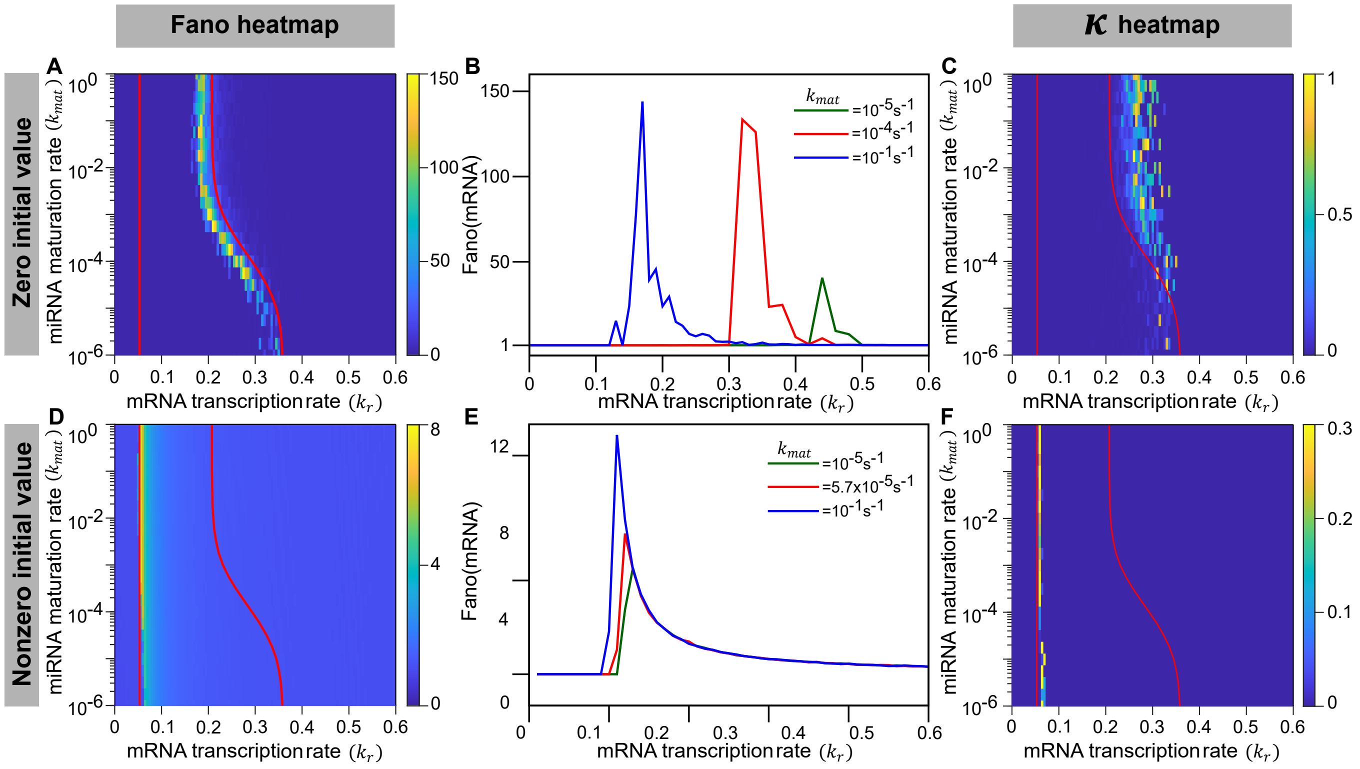


Figure S1: **(Related to Fig. 4 in main text) Initial condition-dependent mRNA noise and bimodality near bifurcation boundaries in positive feedback for** $(g_s/g_m) < (\gamma_s/\gamma_m)$. Results are shown from stochastic simulations with two distinct initial conditions: Zero (panels A-C), and high nonzero (panels D-F) initial copy numbers i.e., $(r_0, p_0, s_0, m_0,) = (0, 0, 0, 0)$ and $(r_0, p_0, s_0, m_0,) = (2000, 2000, 2000, 2000)$, respectively. (A, D) Heatmaps of the steady-state Fano factor of mRNAs are shown in the $k_r$-$k_{mat}$ plane, for zero (A) and nonzero (D) initial conditions. Red solid curves represent the saddle-node bifurcation boundaries obtained from stability analysis of mean-field equations (sections 5 - 6 in SI). (B, E) The steady-state Fano factors of mRNA are plotted as functions of $k_r$ for different maturation rates ($k_{mat}$). Note that the peak positions in the Fano factor shift with the $k_{mat}$, and the peak heights (i.e., maximum noise level) increase with $k_{mat}$. (C, F) The heatmaps of bimodality strength ($\kappa$) are shown in the $k_r$ vs $k_{mat}$ plane. The same red bifurcation curves, as in A and D, are replotted in C and F. Note that high mRNA noise and nonzero bimodality strength ($\kappa$), corresponding to zero and nonzero initial conditions, appear near the right (A and C) and left bifurcation boundaries (D and F), respectively. For all panels, $k_b = 10^{-3}s^{-1}, k_u = 10^{-2}s^{-1}, k_s^0 = 0.5s^{-1}, k_s = 0.05s^{-1}, g_s = 0.000017s^{-1}, g_m = 0.00017s^{-1}, \gamma_s = \gamma_m = 0.04s^{-1}$. Other parameters are from Table I.

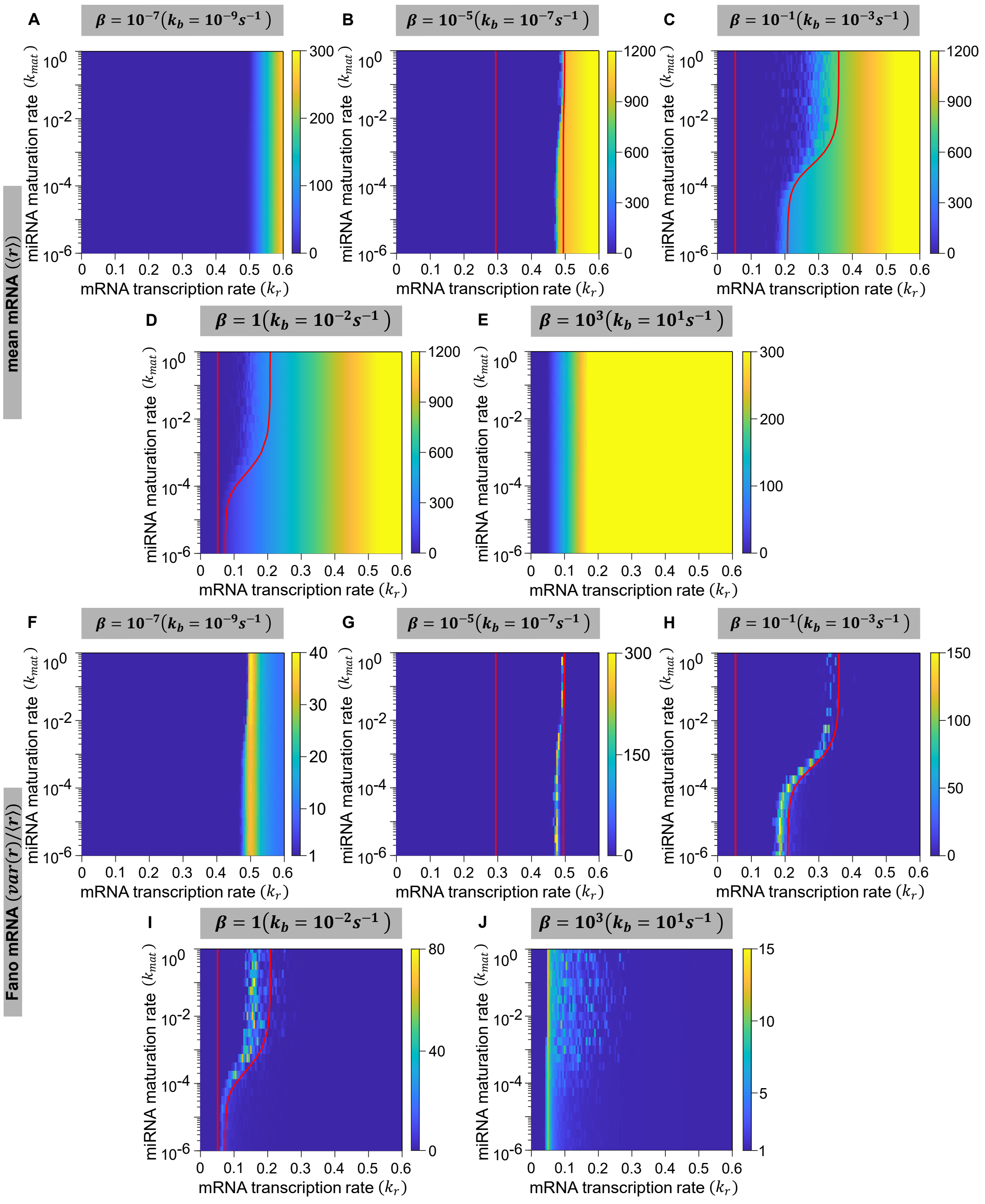


Figure S2: **(Related to Fig. 5 in main text) Effect of varying feedback strength on the mean mRNA and mRNA Fano factor in a miRNA-mediated positive feedback loop.** Heatmaps of mean mRNA level (A-E) and mRNA Fano factor (F-J) in the steady state obtained from stochastic simulations with zero initial condition (i.e., $(r_0, p_0, s_0, m_0) = (0, 0, 0, 0)$). Red solid curves in mean mRNA heatmaps (B, C, D) and Fano factor heatmaps (G, H, I) represent saddle-node bifurcation boundaries, replotted from Fig.5B-D (see main text), corresponding to different feedback strengths ($\beta = 10^{-5}, 10^{-1}, 1$). Note that for all feedback strengths, steady-state mean mRNA undergoes a transition from low to high values, and mRNA noise amplifies around the transition region. Since the initial condition is zero, for intermediate feedback strengths ($\beta = 10^{-5}, 10^{-1}, 1$), the transition of mean mRNA (see B, C, D) and the amplification of mRNA noise (see G, H, I) occur near the right bifurcation boundary (as seen in Fig. 2 and in Fig. 4A of main text). For all panels, $k_u = 10^{-2}s^{-1}, k_s^0 = 0.5s^{-1}, k_s = 0.05s^{-1}, g_s = 0.00017s^{-1}, g_m = 0.000017s^{-1}, \gamma_s = \gamma_m = 0.04s^{-1}$. Other parameters are from Table I.

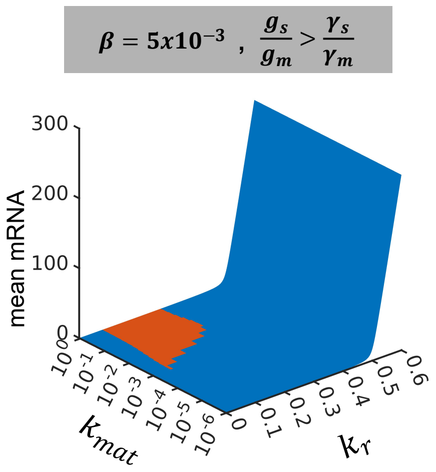


Figure S3: **(Related to Fig. 6B in main text)** The steady-state mean mRNA ($\langle r \rangle$), obtained by numerically solving mean-field equations (Eq. S 3.1-S 3.5), plotted in the $k_r - k_{mat}$ plane. The oscillatory and non-oscillatory regions are indicated in orange and blue, respectively. For all panels, $k_b$ = $5\text{x}10^{-5}s^{-1}, k_u = 10^{-2}s^{-1}(\beta = 5\text{x}10^{-3}), k_s^0 = 0.05s^{-1}, k_s = 0.5s^{-1}, g_s = g_m = 0.000017s^{-1}, \gamma_s = 0.004s^{-1}, \gamma_m = 4s^{-1}$. Other parameter values are taken from Table I (main text).

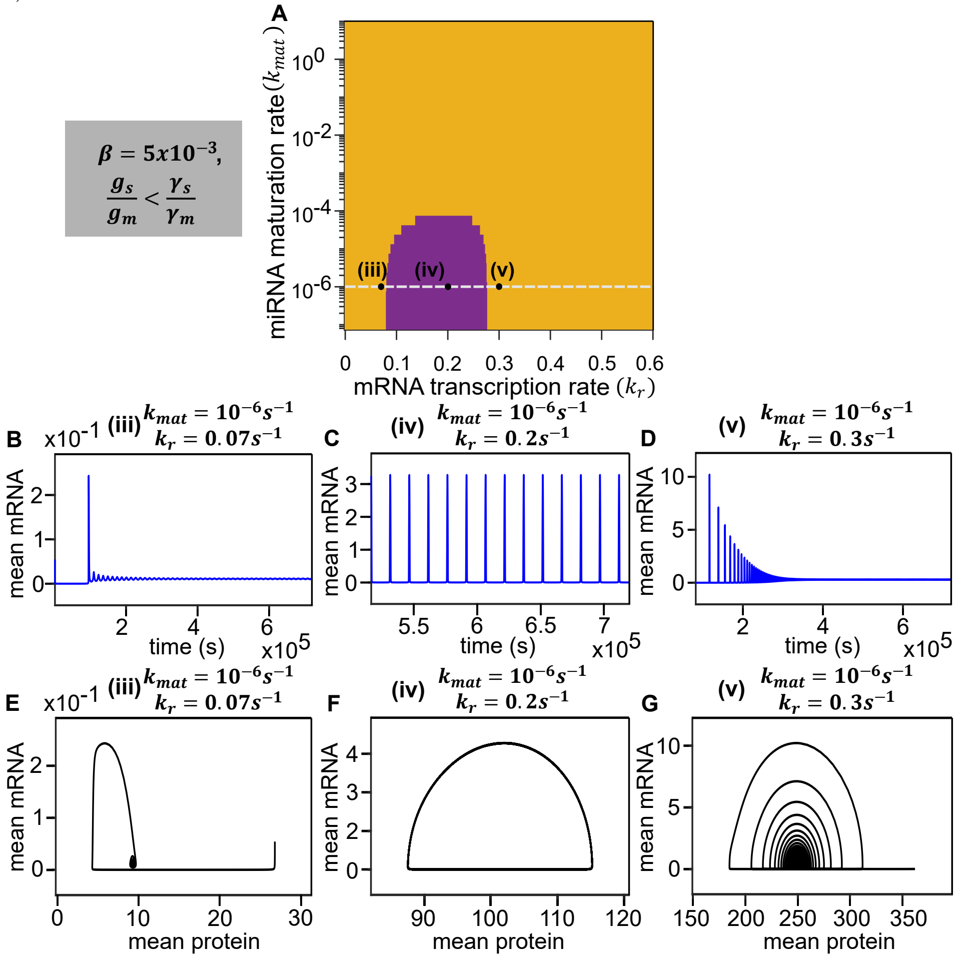


Figure S4: **(Related to Fig. 6D-6H in main text)** (A) The oscillatory (purple) and non-oscillatory (yellow) regions on the $k_r - k_{mat}$ plane, obtained from stability analysis of mean-field equations. The horizontal dashed line represents a fixed value of $k_{mat} = 10^{-6}s^{-1}$ (corresponding to Fig. 6H in main text). The $(k_r, k_{mat})$ coordinates of the highlighted points in A are (along the horizontal dashed line): **(iii)** ($k_r = 0.07s^{-1}, k_{mat} = 10^{-6}s^{-1}$), **(iv)** ($k_r = 0.2s^{-1}, k_{mat} = 10^{-2}s^{-1}$), and **(v)** ($k_r = 0.3s^{-1}, k_{mat} = 10s^{-1}$). (B-D) The steady-state mean mRNA time profiles, obtained from mean-field equations, are shown (corresponding to points **(iii), (iv)** and **(v)**). Note that the mean mRNA exhibits sustained oscillation for point **(iv)** (C), which lies within the oscillatory region. For the other two points, **(iii)** and **(v)**, the mean mRNA shows damped oscillation (B and D). (E-G) Phase portraits in the plane of mean mRNA vs mean protein, corresponding to points **(iii), (iv)** and **(v)** are shown. The phase portrait for the point **(iv)** is a limit cycle (see F), and for points **(iii)** and **(v)**, they are stable spirals (E and G). For all panels, $k_b$ = $5\text{x}10^{-5}s^{-1}, k_u = 10^{-2}s^{-1}(\beta = 5\text{x}10^{-3}), k_s^0 = 0.05s^{-1}, k_s = 0.5s^{-1}, g_s = g_m = 0.000017s^{-1}, \gamma_s = 4s^{-1}, \gamma_m = 0.004s^{-1}$. Other parameter values are taken from Table I (main text).

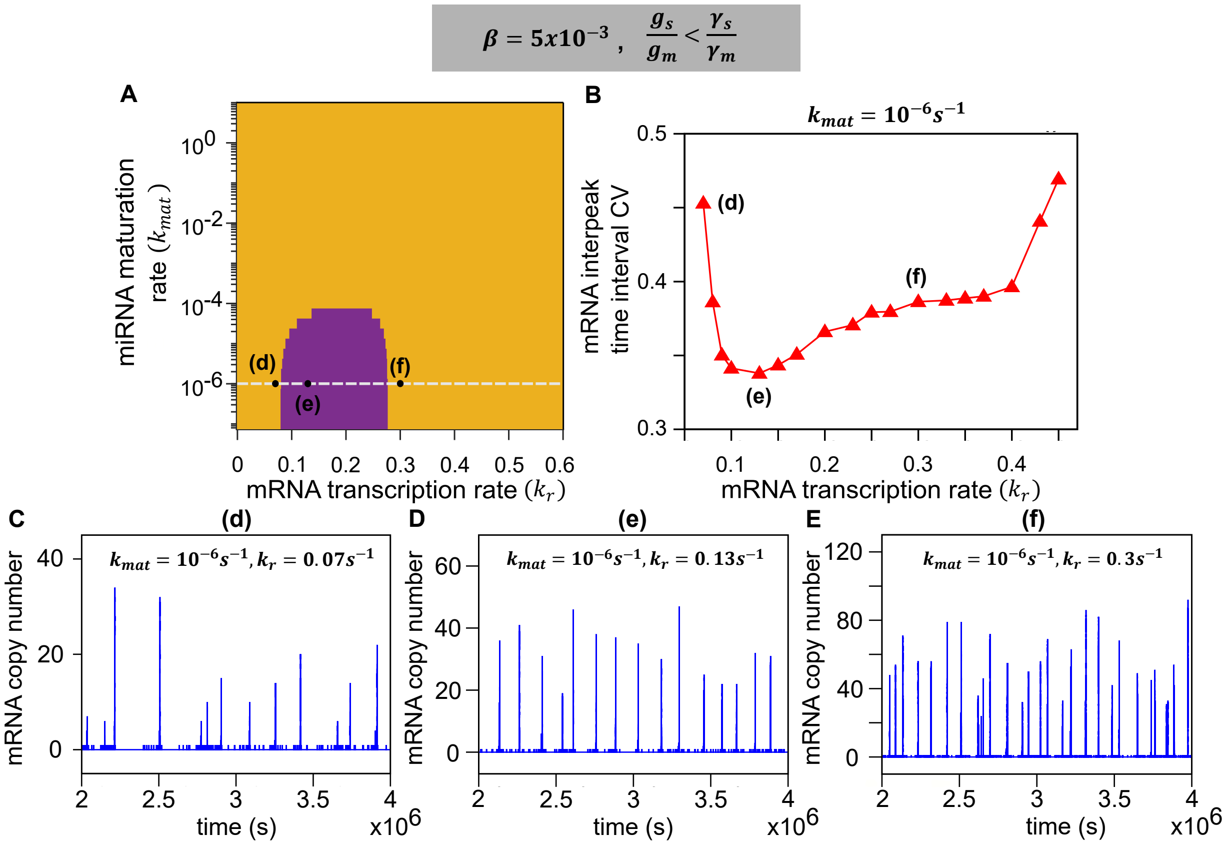


Figure S5: **(Related to Fig. 6L-6O in main text)** (A) The oscillatory (purple) and non-oscillatory (yellow) regions on the $k_r - k_{mat}$ plane, obtained from stability analysis of mean-field equations. The horizontal dashed line represents a fixed value of $k_{mat} = 10^{-6}s^{-1}$. The $(k_r, k_{mat})$ coordinates of the points positioned on the horizontal dashed line in A are: **(d)** ($k_r = 0.07s^{-1}, k_{mat} = 10^{-6}s^{-1}$), **(e)** ($k_r = 0.13s^{-1}, k_{mat} = 10^{-6}s^{-1}$), and **(f)** ($k_r = 0.3s^{-1}, k_{mat} = 10^{-6}s^{-1}$). (B) For a fixed $k_{mat} = 10^{-6}s^{-1}$ (along the horizontal dashed line in A), the CV of mRNA interpeak time interval, $CV_\tau$, is plotted with varying $k_r$ values. The $CV_\tau$ values show non-monotonic behavior with increasing $k_r$. The $CV_\tau$ values of the points **(d), (e)** and **(f)** are indicated in B. (C-E) The steady-state trajectories of mRNA copy numbers, obtained from stochastic simulations, corresponding to the points **(d)**, **(e)**, and **(f)** are shown. The appearance of peaks in the mRNA trajectories is irregular for **(d)**, then becomes regular for **(e)**, and again becomes irregular for **(f)**. For all panels, $k_b = 5\text{x}10^{-5}s^{-1}, k_u = 10^{-2}s^{-1}(\beta = 5\text{x}10^{-3}), k_s^0 = 0.05s^{-1}, k_s = 0.5s^{-1}, g_s = g_m = 0.000017s^{-1}, \gamma_s = 4s^{-1}, \gamma_m = 0.004s^{-1}$. For (B-E), $k_{mat} = 10^{-6}s^{-1}$. Other parameters are taken from Table I (main text).

### Mathematical Analysis of The Model

### 1 Stochastic Master Equation

In our model, there are four stochastic variables, which are the copy number of mRNAs, proteins, precursor miRNAs and mature miRNAs, denoted by $r(t), p(t), s(t)$ & $m(t)$ respectively. In addition there is a state variable $i(t)$ describing state of the miRNA coding gene at any time $t$. Let $P^u_{r,p,s,m}(t) := P(r,p,s,m,i=0,t)$ and $P^b_{r,p,s,m}(t) := P(r,p,s,m,i=1,t)$ be the probabilities that miRNA coding gene is in *Unbound* or *Bound* state, and there are $r$ number of mRNAs, $p$ number of proteins, $s$ number of precursor miRNAs and $m$ number of mature miRNAs at any time $t$, respectively. Therefore, $P_{r,p,s,m}(t) := P(r,p,s,m,t) = P^u_{r,p,s,m}(t)+P^b_{r,p,s,m}(t)$ be the probability that there are $r$ number of mRNAs, $p$ number of proteins, $s$ number of precursor miRNAs & $m$ number of mature miRNAs at any time $t$, respectively.

The master equation corresponding to *Bound* and *Unbound* states are,

$$\begin{aligned}\frac{\partial P^b_{r,p,s,m}}{\partial t} &= k_r\left[P^b_{r-1,p,s,m}-P^b_{r,p,s,m}\right]+g_r\left[(r+1)P^b_{r+1,p,s,m}-rP^b_{r,p,s,m}\right]\\ &+k_p\left[rP^b_{r,p-1,s,m}-rP^b_{r,p,s,m}\right]+g_p\left[(p+1)P^b_{r,p+1,s,m}-pP^b_{r,p,s,m}\right]\\ &+K_b(p+1)P^u_{r,p+1,s,m}-K_uP^b_{r,p,s,m}\\ &+k_s\left[P^b_{r,p,s-1,m}-P^b_{r,p,s,m}\right]+g_s\left[(s+1)P^b_{r,p,s+1,m}-sP^b_{r,p,s,m}\right]\\ &+K_{mat}\left[(s+1)P^b_{r,p,s+1,m-1}-sP^b_{r,p,s,m}\right]+g_m\left[(m+1)P^b_{r,p,s,m+1}-mP^b_{r,p,s,m}\right]\\ &+\gamma_s\left[(r+1)(s+1)P^b_{r+1,p,s+1,m}-rsP^b_{r,p,s,m}\right]+\gamma_m\left[(r+1)(m+1)P^b_{r+1,p,s,m+1}-rmP^b_{r,p,s,m}\right]\end{aligned} \quad \text{(S 1.1)}$$

and,

$$\begin{aligned}\frac{\partial P^u_{r,p,s,m}}{\partial t} &= k_r\left[P^u_{r-1,p,s,m}-P^u_{r,p,s,m}\right]+g_r\left[(r+1)P^u_{r+1,p,s,m}-rP^u_{r,p,s,m}\right]\\ &+k_p\left[rP^u_{r,p-1,s,m}-rP^u_{r,p,s,m}\right]+g_p\left[(p+1)P^u_{r,p+1,s,m}-pP^u_{r,p,s,m}\right]\\ &+K_uP^b_{r,p-1,s,m}-K_bpP^u_{r,p,s,m}\\ &+k^0_s\left[P^u_{r,p,s-1,m}-P^u_{r,p,s,m}\right]+g_s\left[(s+1)P^u_{r,p,s+1,m}-sP^u_{r,p,s,m}\right]\\ &+K_{mat}\left[(s+1)P^u_{r,p,s+1,m-1}-sP^u_{r,p,s,m}\right]+g_m\left[(m+1)P^u_{r,p,s,m+1}-mP^u_{r,p,s,m}\right]\\ &+\gamma_s\left[(r+1)(s+1)P^u_{r+1,p,s+1,m}-rsP^u_{r,p,s,m}\right]+\gamma_m\left[(r+1)(m+1)P^u_{r+1,p,s,m+1}-rmP^u_{r,p,s,m}\right]\end{aligned} \quad \text{(S 1.2)}$$

Therefore, using S 1.1 & S 1.2, we get the chemical master equation as follows,

$$\begin{aligned}\frac{\partial P_{r,p,s,m}}{\partial t} &= \left[\frac{\partial P^b_{r,p,s,m}}{\partial t}+\frac{\partial P^u_{r,p,s,m}}{\partial t}\right]\\ \implies \frac{\partial P_{r,p,s,m}}{\partial t} &= k_r\left[P_{r-1,p,s,m}-P_{r,p,s,m}\right]+g_r\left[(r+1)P_{r+1,p,s,m}-rP_{r,p,s,m}\right]\\ &+k_p\left[rP_{r,p-1,s,m}-rP_{r,p,s,m}\right]+g_p\left[(p+1)P_{r,p+1,s,m}-pP_{r,p,s,m}\right]\\ &+K_u\left[P^b_{r,p-1,s,m}-P^b_{r,p,s,m}\right]+K_b\left[(p+1)P^u_{r,p+1,s,m}-pP^u_{r,p,s,m}\right]\\ &+k^0_s\left[P^u_{r,p,s-1,m}-P^u_{r,p,s,m}\right]+k_s\left[P^b_{r,p,s-1,m}-P^b_{r,p,s,m}\right]+g_s\left[(s+1)P_{r,p,s+1,m}-sP_{r,p,s,m}\right]\\ &+K_{mat}\left[(s+1)P_{r,p,s+1,m-1}-sP_{r,p,s,m}\right]+g_m\left[(m+1)P_{r,p,s,m+1}-mP_{r,p,s,m}\right]\\ &+\gamma_s\left[(r+1)(s+1)P_{r+1,p,s+1,m}-rsP_{r,p,s,m}\right]+\gamma_m\left[(r+1)(m+1)P_{r+1,p,s,m+1}-rmP_{r,p,s,m}\right]\end{aligned} \quad \text{(S 1.3)}$$

### 2 Equations of first order moments

To explore the miRNA-mediated feedback model analytically at the mean level we derive the first order moments as defined below,

$$\begin{aligned}\langle r\rangle &= \sum_{r,p,s,m=0}^{\infty} rP_{r,p,s,m} = \sum_{r,p,s,m=0}^{\infty} r\left(P^b_{r,p,s,m}+P^u_{r,p,s,m}\right) = \langle r\rangle^b+\langle r\rangle^u,\\ \langle p\rangle &= \sum_{r,p,s,m=0}^{\infty} pP_{r,p,s,m} = \sum_{r,p,s,m=0}^{\infty} p\left(P^b_{r,p,s,m}+P^u_{r,p,s,m}\right) = \langle p\rangle^b+\langle p\rangle^u \text{etc.}\end{aligned}$$

Multiplying equation S 1.3 by $r$ and summing up over all variables, we get,

$$\begin{aligned}
\sum_{r,p,s,m=0}^{\infty} r\frac{\partial P_{r,p,s,m}}{\partial t} &= k_r\left[\sum_{r,p,s,m=0}^{\infty} rP_{r-1,p,s,m} - \sum_{r,p,s,m=0}^{\infty} rP_{r,p,s,m}\right] + g_r\left[\sum_{r,p,s,m=0}^{\infty} r(r+1)P_{r+1,p,s,m} - \sum_{r,p,s,m=0}^{\infty} r^2P_{r,p,s,m}\right]\\
&+ k_p\left[\sum_{r,p,s,m=0}^{\infty} r^2P_{r,p-1,s,m} - \sum_{r,p,s,m=0}^{\infty} r^2P_{r,p,s,m}\right] + g_p\left[\sum_{r,p,s,m=0}^{\infty} r(p+1)P_{r,p+1,s,m} - \sum_{r,p,s,m=0}^{\infty} rpP_{r,p,s,m}\right]\\
&+ K_b\left[\sum_{r,p,s,m=0}^{\infty} r(p+1)P^u_{r,p+1,s,m} - \sum_{r,p,s,m=0}^{\infty} rpP^u_{r,p,s,m}\right] + K_u\left[\sum_{r,p,s,m=0}^{\infty} rP^b_{r,p-1,s,m} - \sum_{r,p,s,m=0}^{\infty} rP^b_{r,p,s,m}\right]\\
&+ k_s\left[\sum_{r,p,s,m=0}^{\infty} rP^b_{r,p,s-1,m} - \sum_{r,p,s,m=0}^{\infty} rP^b_{r,p,s,m}\right] + k_s^0\left[\sum_{r,p,s,m=0}^{\infty} rP^u_{r,p,s-1,m} - \sum_{r,p,s,m=0}^{\infty} rP^u_{r,p,s,m}\right]\\
&+ g_s\left[\sum_{r,p,s,m=0}^{\infty} r(s+1)P_{r,p,s+1,m} - \sum_{r,p,s,m=0}^{\infty} rsP_{r,p,s,m}\right]\\
&+ K_{mat}\left[\sum_{r,p,s,m=0}^{\infty} r(s+1)P_{r+1,p,s+1,m} - \sum_{r,p,s,m=0}^{\infty} rsP_{r,p,s,m}\right]\\
&+ g_m\left[\sum_{r,p,s,m=0}^{\infty} r(m+1)P_{r,p,s,m+1} - \sum_{r,p,s,m=0}^{\infty} rmP_{r,p,s,m}\right]\\
&+ \gamma_s\left[\sum_{r,p,s,m=0}^{\infty} r(r+1)(s+1)P_{r+1,p,s+1,m} - \sum_{r,p,s,m=0}^{\infty} r^2sP_{r,p,s,m}\right]\\
&+ \gamma_m\left[\sum_{r,p,s,m=0}^{\infty} r(r+1)(m+1)P_{r+1,p,s,m+1} - \sum_{r,p,s,m=0}^{\infty} r^2mP_{r,p,s,m}\right]\\
\Longrightarrow \frac{\partial\langle r\rangle}{\partial t} &= k_r\left[\sum_{r,p,s,m=0}^{\infty} (r+1)P_{r,p,s,m} - \sum_{r,p,s,m=0}^{\infty} rP_{r,p,s,m}\right] + g_r\left[\sum_{r,p,s,m=0}^{\infty} (r-1)rP_{r,p,s,m} - \sum_{r,p,s,m=0}^{\infty} r^2P_{r,p,s,m}\right]\\
&+ \gamma_s\left[\sum_{r,p,s,m=0}^{\infty} (r-1)rsP_{r,p,s,m} - \sum_{r,p,s,m=0}^{\infty} r^2sP_{r,p,s,m}\right]\\
&+ \gamma_m\left[\sum_{r,p,s,m=0}^{\infty} (r-1)rmP_{r,p,s,m} - \sum_{r,p,s,m=0}^{\infty} r^2mP_{r,p,s,m}\right]
\end{aligned}$$

This simplifies to

- $$\frac{\partial\langle r\rangle}{\partial t} = k_r - g_r\langle r\rangle - \gamma_s\langle rs\rangle + \gamma_m\langle rm\rangle \tag{S 2.1}$$

Using similar technique, we derive the time-evolution equation for all the first order moments as below,

- $$\frac{\partial\langle p\rangle}{\partial t} = k_p\langle r\rangle - g_p\langle p\rangle + k_uP^b - k_b\langle p\rangle^u \tag{S 2.2}$$
- $$\frac{\partial\langle s\rangle}{\partial t} = k_s^0P^u + k_sP^b - g_s\langle s\rangle - k_{mat}\langle s\rangle - \gamma_s\langle rs\rangle \tag{S 2.3}$$
- $$\frac{\partial\langle m\rangle}{\partial t} = k_{mat}\langle s\rangle - g_m\langle m\rangle - \gamma_m\langle rm\rangle \tag{S 2.4}$$

where, $P^b := \sum_{r,p,s,m=0}^{\infty} P^b_{r,p,s,m}$ be the probability that miRNA coding gene is in 'Bound' state, and $P^u := \sum_{r,p,s,m=0}^{\infty} P^u_{r,p,s,m}$ be the probability that miRNA coding gene is in 'Unbound' state. Thus $P^b + P^u = 1$.

Now form S 1.1 summing up in $r, p, s$ & $m$ we get,

$$
\begin{aligned}
\sum_{r,p,s,m=0}^{\infty} \frac{\partial P^b_{r,p,s,m}}{\partial t} &= k_r \left[ \sum_{r,p,s,m=0}^{\infty} P^b_{r-1,p,s,m} - \sum_{r,p,s,m=0}^{\infty} P^b_{r,p,s,m} \right] + g_r \left[ \sum_{r,p,s,m=0}^{\infty} (r+1) P^b_{r+1,p,s,m} - \sum_{r,p,s,m=0}^{\infty} r P^b_{r,p,s,m} \right] \\
&+ k_p \left[ \sum_{r,p,s,m=0}^{\infty} r P^b_{r,p-1,s,m} - \sum_{r,p,s,m=0}^{\infty} r P^b_{r,p,s,m} \right] + g_p \left[ \sum_{r,p,s,m=0}^{\infty} (p+1) P^b_{r,p+1,s,m} - \sum_{r,p,s,m=0}^{\infty} p P^b_{r,p,s,m} \right] \\
&+ K_b \sum_{r,p,s,m=0}^{\infty} (p+1) P^u_{r,p+1,s,m} - K_u \sum_{r,p,s,m=0}^{\infty} P^b_{r,p,s,m} + k_s \left[ \sum_{r,p,s,m=0}^{\infty} P^b_{r,p,s-1,m} - \sum_{r,p,s,m=0}^{\infty} P^b_{r,p,s,m} \right] \\
&+ g_s \left[ \sum_{r,p,s,m=0}^{\infty} (s+1) P^b_{r,p,s+1,m} - \sum_{r,p,s,m=0}^{\infty} s P^b_{r,p,s,m} \right] \\
&+ K_{mat} \left[ \sum_{r,p,s,m=0}^{\infty} (s+1) P^b_{r,p,s+1,m-1} - \sum_{r,p,s,m=0}^{\infty} s P^b_{r,p,s,m} \right] \\
&+ g_m \left[ \sum_{r,p,s,m=0}^{\infty} (m+1) P^b_{r,p,s,m+1} - \sum_{r,p,s,m=0}^{\infty} m P^b_{r,p,s,m} \right] \\
&+ \gamma_s \left[ \sum_{r,p,s,m=0}^{\infty} (r+1)(s+1) P^b_{r+1,p,s+1,m} - \sum_{r,p,s,m=0}^{\infty} rs P^b_{r,p,s,m} \right] \\
&+ \gamma_m \left[ \sum_{r,p,s,m=0}^{\infty} (r+1)(m+1) P^b_{r+1,p,s,m+1} - \sum_{r,p,s,m=0}^{\infty} rm P^b_{r,p,s,m} \right] \\
\implies \frac{\partial P^b}{\partial t} &= k_b \langle p \rangle^u - k_u P^b \qquad \text{(S 2.5)}
\end{aligned}
$$

Equation S 2.5 describes the time-evolution of the probability of miRNA coding gene being in the 'Bound' state.

Note that the moment equations of $\langle r \rangle, \langle s \rangle$ & $\langle m \rangle$ contain the next second-order moments, creating an infinite hierarchy. Therefore, the equations S 2.1 - S 2.5 do not form a closed system. We assume a mean-field approximation (**MFA**) to break the hierarchy, where $2nd$ order moments are expressed in terms of $1st$ order moments.

### 3 Mean-Field Approximation

First order moment (**FOM**) of variables $r(t), p(t), s(t)$ & $c(t)$ describe the average copy number of mRNAs, proteins, precursor miRNAs & mature miRNAs, respectively, at time t. From equations S 2.1 - S 2.5 we have the following,

$$
\begin{aligned}
\textbf{FOM 1}: \frac{\partial \langle r \rangle}{\partial t} &= k_r - g_r \langle r \rangle - \gamma_s \langle rs \rangle + \gamma_m \langle rm \rangle \\
\textbf{FOM 2}: \frac{\partial \langle p \rangle}{\partial t} &= k_p \langle r \rangle - g_p \langle p \rangle + k_u P^b - k_b \langle p \rangle^u \\
\textbf{FOM 3}: \frac{\partial \langle s \rangle}{\partial t} &= \left[ k_s P^b + k_s^0 (1 - P^b) \right] - g_s \langle s \rangle - k_{mat} \langle s \rangle - \gamma_s \langle rs \rangle \\
\textbf{FOM 4}: \frac{\partial \langle m \rangle}{\partial t} &= k_{mat} \langle s \rangle - g_m \langle m \rangle - \gamma_s \langle rm \rangle \\
\textbf{FOM 5}: \frac{\partial P^b}{\partial t} &= k_b \langle p \rangle^u - k_u P^b \qquad \left[ where, P^b + P^u = 1 \right]
\end{aligned}
$$

which describe the time evolution of mean mRNA($\langle r \rangle$), protein($\langle p \rangle$), precursor miRNA($\langle s \rangle$) and mature miRNA ($\langle m \rangle$) copy number. The equation **FOM 5** is to be considered along with **FOM 1 - 4** in order to solve for $\langle r \rangle, \langle p \rangle, \langle s \rangle$ & $\langle m \rangle$, since $P^b$ is present in **FOM 2** and **FOM 3**

The second order moments in **FOM 1 - 5** are $\langle rs \rangle$ & $\langle rm \rangle$. Also, $\langle p \rangle^u$ is another variable that breaks the closure of the system of equations **FOM 1 - 5**. So we define the mean-field approximations (**MFA**) as below to express $\langle rs \rangle, \langle rm \rangle$ and $\langle p \rangle^u$ in terms of first-order moments.

$$
\begin{aligned}
&\textbf{MFA I.} \langle (r - \langle r \rangle)(s - \langle s \rangle) \rangle = 0 \implies \langle rs \rangle = \langle r \rangle \langle s \rangle \\
&\textbf{MFA II.} \langle (r - \langle r \rangle)(m - \langle m \rangle) \rangle = 0 \implies \langle rm \rangle = \langle r \rangle \langle m \rangle \\
&\textbf{MFA III.} \langle (p - \langle p \rangle) \rangle^u = 0 \implies \langle p \rangle^u = \langle p \rangle P^u = \langle p \rangle \left(1 - P^b\right)
\end{aligned}
$$

we get from **FOM 1 - 5** the following,

$$\frac{\partial\langle r\rangle}{\partial t} \approx k_r - g_r\langle r\rangle - \gamma_s\langle r\rangle\langle s\rangle - \gamma_m\langle r\rangle\langle m\rangle \tag{S 3.1}$$

$$\frac{\partial\langle p\rangle}{\partial t} \approx k_p\langle r\rangle - g_p\langle p\rangle + k_u P^b - k_b\langle p\rangle(1-P^b) \tag{S 3.2}$$

$$\frac{\partial\langle s\rangle}{\partial t} \approx \left[k_s P^b + k_s^0(1-P^b)\right] - g_s\langle s\rangle - k_{mat}\langle s\rangle - \gamma_s\langle r\rangle\langle s\rangle \tag{S 3.3}$$

$$\frac{\partial\langle m\rangle}{\partial t} \approx k_{mat}\langle s\rangle - g_m\langle m\rangle - \gamma_m\langle r\rangle\langle m\rangle \tag{S 3.4}$$

$$\frac{\partial P^b}{\partial t} \approx k_b\langle p\rangle\left(1-P^b\right) - k_u P^b \tag{S 3.5}$$

which we shall refer to as Mean-Field Equations (**MFE**).

### 4 Steady state solution of Mean-Field Approximated Equations

From S 3.2 & S 3.5 at steady state we have,

$$k_p\langle r\rangle - g_p\langle p\rangle + k_u P^b - k_b\langle p\rangle(1-P^b) = 0$$
$$k_b\langle p\rangle\left(1-P^b\right) - k_u P^b = 0$$

Therefore,

$$\langle r\rangle k_p - g_p\langle p\rangle = 0$$
$$\implies \langle p\rangle = \frac{k_p}{g_p}\langle r\rangle \tag{S 4.1}$$

Now,

$$P^b = \frac{k_b\langle p\rangle}{(k_b\langle p\rangle + k_u)}$$
$$\implies P^b = \frac{k_p k_b\langle r\rangle}{(k_p k_b\langle r\rangle + g_p k_u)} \tag{S 4.2}$$

From S 3.3 in steady state,

$$k_s^0(1-P^b) + k_s P^b - g_s\langle s\rangle - k_{mat}\langle s\rangle - \gamma_s\langle r\rangle\langle s\rangle = 0$$
$$\implies \langle s\rangle = -\frac{k_s^0 P^b - k_s^0 - k_s P^b}{g_s + k_{mat} + \gamma_s\langle s\rangle}$$

From S 3.4 in steady state,

$$k_{mat}\langle s\rangle - g_m\langle m\rangle - \gamma_m\langle r\rangle\langle m\rangle = 0$$
$$\implies \langle m\rangle = \frac{k_{mat}\langle s\rangle}{g_m + \gamma_m\langle r\rangle}$$
$$\implies \langle m\rangle = -\frac{k_{mat}\left(k_s^0 P^b - k_s^0 - k_s P^b\right)}{(g_m + \gamma_m\langle r\rangle)\left(g_s + k_{mat} + \gamma_s\langle s\rangle\right)}$$

Using S 4.1 & S 4.2 in above equations we get,

$$\langle s\rangle = \frac{\left(g_p k_s^0 k_u + k_b k_p k_s\langle r\rangle\right)}{(g_p k_u + k_b k_p\langle r\rangle)\left(g_s + k_{mat} + \gamma_s\langle r\rangle\right)} \tag{S 4.3}$$

$$\langle m\rangle = \frac{k_{mat}\left(g_p k_s^0 k_u + k_b k_p k_s\langle r\rangle\right)}{(g_m + \langle r\rangle\gamma_m)\left(g_p k_u + k_b k_p\langle r\rangle\right)\left(g_s + k_{mat} + \gamma_s\langle r\rangle\right)} \tag{S 4.4}$$

Now, from S 3.1 in the steady state we get,

$$k_r - g_r\langle r\rangle - \gamma_s\langle r\rangle\langle s\rangle + \gamma_m\langle r\rangle\langle m\rangle$$
$$\langle r\rangle = \frac{k_r}{g_r + \gamma_s\langle s\rangle + \gamma_m\langle m\rangle}$$

Substituting S 4.3 & S 4.4 in the above equation and then simplifying, we get the following polynomial equation of $\langle r\rangle$,

$$a_4\langle r\rangle^4 + a_3\langle r\rangle^3 + a_2\langle r\rangle^2 + a_1\langle r\rangle + a_0 = 0 \tag{S 4.5}$$

where

$$a_4 = g_r k_b k_p \gamma_s \gamma_m \tag{S 4.6}$$

$$a_3 = g_r k_b (g_s + k_{mat}) k_p \gamma_m + k_b k_p (k_s - k_r) \gamma_s \gamma_m + g_r (g_m k_b k_p + g_p k_u \gamma_m) \gamma_s \tag{S 4.7}$$

$$\begin{aligned} a_2 &= g_m k_b k_p (k_s - k_r) \gamma_s + g_m g_r \{k_b (g_s + k_{mat}) k_p + g_p k_u \gamma_s\} \\ &+ \gamma_m \Big[ k_b k_{mat} k_p (k_s - k_r) + g_s (g_p g_r k_u - k_b k_p k_r) + g_p k_u \left( g_r k_{mat} - k_r \gamma_s + k_s^0 \gamma_s \right) \Big] \end{aligned} \tag{S 4.8}$$

$$\begin{aligned} a_1 &= -g_m (g_s + k_{mat}) (k_b k_p k_r - g_p g_r k_u) - g_p (g_s + k_{mat}) k_r k_u \gamma_m \\ &+ g_p k_{mat} k_s^0 k_u \gamma_m + g_m g_p \left( k_s^0 - k_r \right) k_u \gamma_s \end{aligned} \tag{S 4.9}$$

$$a_0 = -g_m g_p (g_s + k_{mat}) k_r k_u \tag{S 4.10}$$

Equation S 4.5 can also be expressed as,

$$\langle r \rangle^4 + b_3 \langle r \rangle^3 + b_2 \langle r \rangle^2 + b_1 \langle r \rangle + b_0 = 0 \tag{S 4.11}$$

where,

$$b_i = \frac{a_i}{a_4} \forall i = 0, 1, 2, 3 \text{ provided } a_4 \neq 0 \tag{S 4.12}$$

### 5 Multi-stability of sytsem

Let $\Delta$ be the discriminant of Eq. S 4.5 (equivalently S 4.11). From the properties of the discriminant of a $nth$ degree polynomial equation, we know that

- polynomial has multiple roots if and only if $\Delta = 0$.
- if $\Delta > 0$, then there is an integer $k \leq \frac{n}{4}$, such that there are $2k$ pair of complex conjugate roots, and, $(n - 2 \cdot 2k)$ numbers of real roots.
- if $\Delta < 0$, then there is an integer $k \leq \frac{(n-2)}{4}$, such that there are $(2k+1)$ pair of complex conjugate roots, and, $(n - 2 \cdot (2k+1))$ numbers of real roots.

In our case, for Eq. S 4.5 (equivalently S 4.11) $n = 4$. Therefore,

**i** we have multiple roots if and only if $\Delta = 0$.

**ii** If $\Delta > 0$, then there is an integer $k \leq 1$, such that there are $2k$ pair of complex conjugate roots, and, $(n - 2 \cdot 2k)$ numbers of real roots of .

  (a) If $k = 0$, then there are no complex conjugate roots, and, all 4 roots are real.

  (b) If $k = 1$, then there is 1 pair of complex conjugate roots, and the remaining 2 roots are real.

**iii** If $\Delta < 0$, then there is an integer $k \leq \frac{1}{2}$, such that there are $(2k+1)$ pairs of complex conjugate roots, and, $(n - 2 \cdot 2k)$ numbers of real roots. Thus $k = 0$, then there is 1 pair of complex conjugate roots, and, remaining 2 roots are real.

*Lemma. The polynomial equation* $\langle r \rangle^4 + b_3 \langle r \rangle^3 + b_2 \langle r \rangle^2 + b_1 \langle r \rangle + b_0 = 0$ *has all it's roots positive iff* $\{b_3 < 0, b_2 > 0, b_1 < 0$ *&* $b_0 > 0\}$.

*proof.* Let $r_1, r_2, r_3, r_4$ are the roots of the $4th$ degree polynomial equation. Then, by vietta's identity,

$$\begin{aligned} &r_1 + r_2 + r_3 + r_4 = -b_3 \\ &r_1 r_2 + r_1 r_3 + r_1 r_4 + r_2 r_3 + r_2 r_4 + r_3 r_4 = b_2 \\ &r_1 r_2 r_3 + r_1 r_2 r_4 + r_1 r_3 r_4 + r_2 r_3 r_4 = -b_1 \\ &r_1 r_2 r_3 r_4 = b_0 \end{aligned}$$

Consider that all the roots are positive. Then using the results from Vieta's formulae,

$$\begin{aligned} &r_1 + r_2 + r_3 + r_4 > 0 \implies -b_3 > 0 \implies b_3 < 0 \\ &r_1 r_2 + r_1 r_3 + r_1 r_4 + r_2 r_3 + r_2 r_4 + r_3 r_4 > 0 \implies b_2 > 0 \\ &r_1 r_2 r_3 + r_1 r_2 r_4 + r_1 r_3 r_4 + r_2 r_3 r_4 > 0 \implies -b_1 > 0 \implies b_1 < 0 \\ &r_1 r_2 r_3 r_4 > 0 \implies b_0 > 0 \end{aligned}$$

Conversely, consider $\{b_3 < 0, b_2 > 0, b_1 < 0 \ \& \ b_0 > 0\}$. Without loss of generality, let $r_1 < 0$. Then from Vieta's formula,

$$b_0 > 0 \implies r_1 r_2 r_3 r_4 > 0 \implies r_2 r_3 r_4 < 0 \qquad [\because r_1 < 0]$$

Now, again using other Vieta's formulae, we get

$$\begin{aligned} b_1 < 0 &\implies -b_1 = r_1r_2r_3 + r_1r_2r_4 + r_1r_3r_4 + r_2r_3r_4 > 0 \\ &\implies r_1\left(r_2r_3 + r_2r_4 + r_3r_4\right) + r_2r_3r_4 > 0 \end{aligned}$$

and,

$$\begin{aligned} b_2 > 0 &\implies r_1r_2 + r_1r_3 + r_1r_4 + r_2r_3 + r_2r_4 + r_3r_4 > 0 \\ &\implies r_1\left(r_2 + r_3 + r_4\right) + r_2r_3 + r_2r_4 + r_3r_4 > 0 \end{aligned}$$

and finally,

$$\begin{aligned} &b_3 < 0 \implies -b_3 = r_1 + r_2 + r_3 + r_4 > 0 \\ &r_2 + r_3 + r_4 > -r_1 > 0 \quad [\because r_1 < 0] \end{aligned}$$

Therefore,

$$r_2r_3 + r_2r_4 + r_3r_4 > -r_1\left(r_2 + r_3 + r_4\right) > 0 \qquad [\because r_1 < 0, r_2 + r_3 + r_4 > 0]$$

and

$$r_2r_3r_4 > -r_1\left(r_2r_3 + r_2r_4 + r_3r_4\right) > 0 \qquad [\because r_1 < 0, r_2r_3 + r_2r_4 + r_3r_4 > 0]$$

This is a contradiction. Therefore all the roots of the polynomial are positive.

For Eq. S 4.5 (equivalently Eq. S 4.11) we have

$$\frac{a_0}{a_4} = b_0 < 0 \qquad [\because a_4 > 0, a_0 < 0, \quad \text{assuming all the parameters of the system are positive.}]$$

Therefore, Eq. S 4.5 (equivalently Eq. S 4.11) cannot have all it's real roots positive.

**i** if $\Delta < 0$, then $k = 0$ and among the 2 real roots at least 1 should be negative.

**ii** if $\Delta > 0$, then either $k = 0$ or $k = 1$.

(a) if $k = 0$, then among the 4 real roots, at least one should be negative.

(b) if $k = 1$, then among the 2 real roots, at least one should be negative.

Hence, the only way multistability can be achieved is if $\Delta > 0$ and $k = 0$, then there is a possibility that among the 4 real roots, only 1 is real & negative and the rest are real & positive. If we achieve this scenario, then only we shall have 2 real & positive roots as 'stable' steady state and the other real & positive root as 'unstable' steady state.

### 6 Stability analysis of the Mean-Field Equations

The Jacobian of the Mean-Field Equations (S 3.1-S 3.5) is,

$$J = \begin{bmatrix} -g_r - m\gamma_m - s\gamma & 0 & -r\gamma_s & -r\gamma_m & 0 \\ k_p & -g_p + k_b(p^b - 1) & 0 & 0 & k_u + k_b p \\ -s\gamma_s & 0 & -g_s - k_{mat} - r\gamma_s & 0 & k_s - k_s^0 \\ -m\gamma_m & 0 & k_{mat} & -g_m - r\gamma_m & 0 \\ 0 & k_b(1 - P^b) & 0 & 0 & -k_u - k_b p \end{bmatrix} \tag{S 6.1}$$

Let $\tilde{X} = (\tilde{r}, \tilde{p}, \tilde{s}, \tilde{m}, \tilde{P^b})$ be a steady state solution of **MFE** (Eq. S 3.1-S 3.5). Then at the steady-state solution $\tilde{X}$ the Jacobian is

$$\begin{aligned} \tilde{J} &= J|_{\tilde{X}=(\tilde{r},\tilde{p},\tilde{s},\tilde{m},\tilde{P^b})} \\ &= \begin{bmatrix} -g_r - \tilde{m}\gamma_m - \tilde{s}\gamma & 0 & -\tilde{r}\gamma_s & -\tilde{r}\gamma_m & 0 \\ k_p & -g_p + k_b(\tilde{P^b} - 1) & 0 & 0 & k_u + k_b\tilde{p} \\ -\tilde{s}\gamma_s & 0 & -g_s - k_{mat} - \tilde{r}\gamma_s & 0 & k_s - k_s^0 \\ -\tilde{m}\gamma_m & 0 & k_{mat} & -g_m - \tilde{r}\gamma_m & 0 \\ 0 & k_b(1 - \tilde{P^b}) & 0 & 0 & -k_u - k_b\tilde{p} \end{bmatrix} \end{aligned} \tag{S 6.2}$$

The eigen values of the Jacobian $\tilde{J}$ is given by

$$|\tilde{J} - \lambda I_5| = 0$$
$$\implies \lambda^5 + c_4\lambda^4 + c_3\lambda^3 + c_2\lambda^2 + c_1\lambda + c_0 = 0 \tag{S 6.3}$$

where,

$$c_4 = g_m + g_p + g_r + g_s + k_{mat} + k_u + k_b(1 + \tilde{p} - \tilde{P^b}) + (\tilde{m} + \tilde{r})\gamma_m + (\tilde{r} + \tilde{s})\gamma_s \tag{S 6.4}$$

$$\begin{aligned}
c_3 = {} & g_r g_s + g_r k_b + g_s k_b + g_r k_{mat} + k_b k_{mat} + g_r k_u + g_s k_u + k_{mat} k_u + g_r k_b \tilde{p} + g_s k_b \tilde{p} + k_b k_{mat} \tilde{p} \\
& - g_r k_b \tilde{P^b} - g_s k_b \tilde{P^b} - k_b k_{mat} \tilde{P^b} + g_s \tilde{m}\gamma_m + k_b \tilde{m}\gamma_m + k_{mat}\tilde{m}\gamma_m + k_u \tilde{m}\gamma_m + k_b \tilde{m}\tilde{p}\gamma_m \\
& - k_b \tilde{m}\tilde{P^b}\gamma_m + g_r \tilde{r}\gamma_m + g_s \tilde{r}\gamma_m + k_b \tilde{r}\gamma_m + k_{mat}\tilde{r}\gamma_m + k_u \tilde{r}\gamma_m + k_b \tilde{p}\tilde{r}\gamma_m - k_b P^b \tilde{r}\gamma_m \\
& + \gamma_s \left[g_r \tilde{r} + (g_s + k_{mat})\tilde{s} + k_u(\tilde{r} + \tilde{s}) + k_b(1 + \tilde{p} - \tilde{P^b})(\tilde{r} + \tilde{s}) + \tilde{r}(\tilde{m} + \tilde{r} + \tilde{s})\gamma_m\right] \\
& + g_m \left[g_p + g_r + g_s + k_b + k_{mat} + k_u + k_b \tilde{p} - k_b \tilde{P^b} + \tilde{m}\gamma_m + (\tilde{r} + \tilde{s})\gamma_s\right] \\
& + g_p \left[g_r + g_s + k_{mat} + k_u + k_b \tilde{p} + (\tilde{m} + \tilde{r})\gamma_m + (\tilde{r} + \tilde{s})\gamma_s\right]
\end{aligned} \tag{S 6.5}$$

$$\begin{aligned}
c_2 = {} & g_r g_s k_b + g_r k_b k_{mat} + g_r g_s k_u + g_r k_{mat} k_u + g_r g_s k_b \tilde{p} + g_r k_b k_{mat} \tilde{p} - g_r g_s k_b \tilde{P^b} - g_r k_b k_{mat} \tilde{P^b} \\
& + g_s k_b \tilde{m}\gamma_m + k_b k_{mat}\tilde{m}\gamma_m + g_s k_u \tilde{m}\gamma_m + k_{mat} k_u \tilde{m}\gamma_m + g_s k_b \tilde{m}\tilde{p}\gamma_m + k_b k_{mat}\tilde{m}\tilde{p}\gamma_m - g_s k_b \tilde{m}\tilde{P^b}\gamma_m \\
& - k_b k_{mat}\tilde{m}\tilde{P^b}\gamma_m + g_r g_s \tilde{r}\gamma_m + g_r k_b \tilde{r}\gamma_m + g_s k_b \tilde{r}\gamma_m + g_r k_{mat}\tilde{r}\gamma_m + k_b k_{mat}\tilde{r}\gamma_m + g_r k_u \tilde{r}\gamma_m + g_s k_u \tilde{r}\gamma_m \\
& + k_{mat} k_u \tilde{r}\gamma_m + g_r k_b \tilde{p}\tilde{r}\gamma_m + g_s k_b \tilde{p}\tilde{r}\gamma_m + k_b k_{mat}\tilde{p}\tilde{r}\gamma_m - g_r k_b \tilde{P^b}\tilde{r}\gamma_m - g_s k_b \tilde{P^b}\tilde{r}\gamma_m - k_b k_{mat}\tilde{P^b}\tilde{r}\gamma_m \\
& + \left[k_u + k_b(1 + \tilde{p} - \tilde{P}b)\right]\left[g_r \tilde{r} + (g_s + k_{mat})\tilde{s}\right]\gamma_s + \tilde{r}\left[g_r \tilde{r} + g_s \tilde{s} + (k_u + k_b(1 + \tilde{p} - \tilde{P^b}))(\tilde{m} + \tilde{r} + \tilde{s})\right]\gamma_m\gamma_s \\
& + g_m \Bigg[g_s k_b + k_b k_{mat} + g_s k_u + k_{mat} k_u + g_s k_b \tilde{p} + k_b k_{mat}\tilde{p} - g_s k_b \tilde{P^b} - k_b k_{mat}\tilde{P^b} + g_s \tilde{m}\gamma_m + k_b \tilde{m}\gamma_m + k_{mat}\tilde{m}\gamma_m \\
& + k_u \tilde{m}\gamma_m + k_b \tilde{m}\tilde{p}\gamma_m - k_b \tilde{m}\tilde{P^b}\gamma_m + \left((g_s + k_{mat})\tilde{s} + k_u(\tilde{r} + \tilde{s}) + k_b(1 + \tilde{p} - \tilde{P^b})(\tilde{r} + \tilde{s}) + \tilde{m}\tilde{r}\gamma_m\right)\gamma_s \\
& + g_r\left(g_s + k_b + k_{mat} + k_u + k_b\tilde{p} - k_b\tilde{P^b} + \tilde{r}\gamma_s\right) + g_p(g_r + g_s + k_{mat} + k_u + k_b\tilde{p} + \tilde{m}\gamma_m + (\tilde{r} + \tilde{s})\gamma_s))\Bigg] \\
& + g_p \Bigg[k_{mat} k_u + k_b k_{mat}\tilde{p} + k_{mat}\tilde{m}\gamma_m + k_u \tilde{m}\gamma_m + k_b \tilde{m}\tilde{p}\gamma_m + k_{mat}\tilde{r}\gamma_m + k_u \tilde{r}\gamma_m + k_b \tilde{p}\tilde{r}\gamma_m \\
& + (k_{mat}\tilde{s} + (k_u + k_b\tilde{p})(\tilde{r} + \tilde{s}) + \tilde{r}(\tilde{m} + \tilde{r} + \tilde{s})\gamma_m)\gamma_s + g_s(k_u + k_b\tilde{p} + (\tilde{m} + \tilde{r})\gamma_m + \tilde{s}\gamma_s) \\
& + g_r(g_s + k_{mat} + k_u + k_b\tilde{p} + \tilde{r}(\gamma_m + \gamma_s))\Bigg]
\end{aligned} \tag{S 6.6}$$

$$\begin{aligned}
c_1 = {} & g_r(g_s + k_{mat})\left(k_u + k_b(1 + \tilde{p} - \tilde{P^b})\right) r\gamma_m \\
& + \tilde{r}\left(-k_b k_p(ks - k_s^0)(-1 + \tilde{P^b}) + \left(k_u + k_b\left(1 + \tilde{p} - \tilde{P^b}\right)\right)(g_r\tilde{r} + g_s\tilde{s})\gamma_m\right)\gamma_s \\
& + g_p \Bigg[(g_s + k_{mat})(k_u + k_b\tilde{p})(\tilde{m} + \tilde{r})\gamma_m + (g_s + k_{mat})(k_u + k_b\tilde{p})\tilde{s}\gamma_s + \tilde{r}(g_s\tilde{s} + (k_u + k_b\tilde{p})(\tilde{m} + \tilde{r} + \tilde{s}))\gamma_m\gamma_s \\
& + g_r\Big((k_u + k_b\tilde{p})\tilde{r}\gamma_m + g_s(k_u + k_b\tilde{p} + \tilde{r}\gamma_m) + k_{mat}(k_u + k_b\tilde{p} + \tilde{r}\gamma_m) + \tilde{r}(k_u + k_b\tilde{p} + \tilde{r}\gamma_m)\gamma_s\Big)\Bigg] \\
& + g_m \Bigg[\left(k_u + k_b(1 + \tilde{p} - \tilde{P^b})\right)((g_s + k_{mat})(g_r + \tilde{m}\gamma_m) + (g_s + k_{mat})\tilde{s}\gamma_s + \tilde{r}(g_r + \tilde{m}\gamma_m)\gamma_s) + \\
& + g_p\Big\{k_{mat} k_u + k_b k_{mat}\tilde{p} + k_{mat}\tilde{m}\gamma_m + k_u\tilde{m}\gamma_m + k_b\tilde{m}\tilde{p}\gamma_m + (k_{mat}\tilde{s} + (k_u + k_b\tilde{p})(\tilde{r} + \tilde{s}) + \tilde{m}\tilde{r}\gamma_m)\gamma_s \\
& + g_r(g_s + k_{mat} + k_u + k_b\tilde{p} + \tilde{r}\gamma_s) + g_s(k_u + k_b\tilde{p} + \tilde{m}\gamma_m + \tilde{s}\gamma_s)\Big\}\Bigg]
\end{aligned} \tag{S 6.7}$$

and,

$$
\begin{aligned}
c_0 &= g_m g_p(k_u + k_b\tilde{p})\left[(g_s + k_{mat})(g_r + \tilde{m}\gamma_m) + (g_r\tilde{r} + (g_s + k_{mat})\tilde{s} + \tilde{m}\tilde{r}\gamma_m)\,\gamma_s\right] \\
&\quad + g_p k_u \tilde{r}\gamma_m\left[g_s\tilde{s}\gamma_s + g_r(g_s + k_{mat} + \tilde{r}\gamma_s)\right] - g_m k_b k_p(k_s - k_s^0)(-1 + \tilde{P^b})\tilde{r}\gamma_s \\
&\quad + k_b\tilde{r}\gamma_m\left[g_p g_r k_{mat}\tilde{p} - k_{mat}k_p(k_s - k_s^0)(-1 + \tilde{P^b}) - k_p(k_s - k_s^0)(-1 + \tilde{P^b})\tilde{r}\gamma_s + g_p\tilde{p}\,(g_s\tilde{s}\gamma_s + g_r(g_s + \tilde{r}\gamma_s))\right]
\end{aligned} \tag{S 6.8}
$$

### 7 Condition under which the bistable region becomes independent of miRNA maturation rate ($k_{mat}$) in miRNA-mediated positive feedback

From the mean-field equations, using S 3.3 and S 3.4 we get,

$$
\begin{aligned}
&\left[k_s P^b + k_s^0(1 - P^b)\right] - (g_s\langle s\rangle + g_m\langle m\rangle) - \langle r\rangle(\gamma_s\langle s\rangle + \gamma_m\langle m\rangle) = 0 \\
\implies &\langle r\rangle(\gamma_s\langle s\rangle + \gamma_m\langle m\rangle) = \left[k_s P^b + k_s^0(1 - P^b)\right] - (g_s\langle s\rangle + g_m\langle m\rangle)
\end{aligned} \tag{S 7.1}
$$

From S 3.1 we get,

$$
\begin{aligned}
&k_r - g_r\langle r\rangle = \langle r\rangle(\gamma_s\langle s\rangle + \gamma_m\langle m\rangle) \\
\implies &k_r - g_r\langle r\rangle = \left[k_s P^b + k_s^0(1 - P^b)\right] - (g_s\langle s\rangle + g_m\langle m\rangle) \\
\implies &\langle r\rangle = \frac{1}{g_r}\left[k_r - \left\{k_s P^b + k_s^0(1 - P^b)\right\} + (g_s\langle s\rangle + g_m\langle m\rangle)\right]
\end{aligned} \tag{S 7.2}
$$

From S 3.4 and S 3.3 we already have,

$$
\begin{aligned}
\langle s\rangle &= \frac{\left[k_s P^b + k_s^0(1 - P^b)\right]}{(g_s + k_{mat} + \gamma_s\langle r\rangle)} \\
\langle m\rangle &= \frac{k_{mat}\langle s\rangle}{(g_m + \langle r\rangle\gamma_m)} = \frac{k_{mat}\left[k_s P^b + k_s^0(1 - P^b)\right]}{(g_m + \langle r\rangle\gamma_m)\,(g_s + k_{mat} + \gamma_s\langle r\rangle)}
\end{aligned}
$$

Using these expressions we get,

$$
\begin{aligned}
(g_s\langle s\rangle + g_m\langle m\rangle) &= \frac{\{g_s(g_m + \gamma_m\langle r\rangle + k_{mat}g_m)\}\left[k_s P^b + k_s^0(1 - P^b)\right]}{(g_m + \langle r\rangle\gamma_m)\,(g_s + k_{mat} + \gamma_s\langle r\rangle)} \\
&= \frac{\{g_m(g_s + k_{mat}) + g_s\gamma_m\langle r\rangle\}\left[k_s P^b + k_s^0(1 - P^b)\right]}{(g_m + \langle r\rangle\gamma_m)\,(g_s + k_{mat} + \gamma_s\langle r\rangle)}
\end{aligned} \tag{S 7.3}
$$

Now, substituting the value of $(g_s\langle s\rangle + g_m\langle m\rangle)$ from S 7.3 into S 7.2 we get

$$
\begin{aligned}
\langle r\rangle &= \frac{1}{g_r}\left[k_r - \left\{k_s P^b + k_s^0(1 - P^b)\right\} + \frac{\{g_m(g_s + k_{mat}) + g_s\gamma_m\langle r\rangle\}\left[k_s P^b + k_s^0(1 - P^b)\right]}{(g_m + \langle r\rangle\gamma_m)\,(g_s + k_{mat} + \gamma_s\langle r\rangle)}\right] \\
&= \frac{1}{g_r}\left[k_r - \frac{\left\{k_s P^b + k_s^0(1 - P^b)\right\}(g_m\gamma_s\langle r\rangle + k_{mat}\gamma_m\langle r\rangle + \gamma_s\gamma_m\langle r\rangle^2)}{(g_m + \langle r\rangle\gamma_m)\,(g_s + k_{mat} + \gamma_s\langle r\rangle)}\right] \\
&= \frac{1}{g_r}\left[k_r - \frac{\left\{(k_s - k_s^0)P^b + k_s^0\right\}(g_m\gamma_s\langle r\rangle + k_{mat}\gamma_m\langle r\rangle + \gamma_s\gamma_m\langle r\rangle^2)}{(g_m + \langle r\rangle\gamma_m)\,(g_s + k_{mat} + \gamma_s\langle r\rangle)}\right] \\
&= \frac{1}{g_r}\left[k_r - \frac{\left\{k_s k_p k_b\langle r\rangle + k_s^0 g_p k_u\right\}\langle r\rangle(g_m\gamma_s + k_{mat}\gamma_m + \gamma_s\gamma_m\langle r\rangle)}{(g_m + \langle r\rangle\gamma_m)\,(g_s + k_{mat} + \gamma_s\langle r\rangle)\,(k_b k_p\langle r\rangle + g_p k_u)}\right] \quad \text{[using S 4.2]}
\end{aligned} \tag{S 7.4}
$$

Eq. S 7.4 gives a polynomial in $\langle r\rangle$ and solving this we obtain $\langle r\rangle$ as a function of both $k_r$ and $k_{mat}$ along with other parameters $g_r, k_p, g_p, k_u, k_b, k_s^0, k_s, g_s, g_m, \gamma_s, \gamma_m$.

#### 7.1 consider $\frac{g_s}{g_m} = \frac{\gamma_s}{\gamma_m} = \alpha$

From Eq. S 7.3 we get,

$$
\begin{aligned}
(g_s\langle s\rangle + g_m\langle m\rangle) &= \frac{\{g_m(\alpha g_m + k_{mat}) + \alpha g_m\gamma_m\langle r\rangle\}\left[k_s P^b + k_s^0(1 - P^b)\right]}{(g_m + \langle r\rangle\gamma_m)\,(\alpha g_m + k_{mat} + \alpha\gamma_m\langle r\rangle)} \\
&= \frac{g_m\,(\alpha g_m + k_{mat} + \alpha\gamma_m\langle r\rangle)\left[k_s P^b + k_s^0(1 - P^b)\right]}{(g_m + \langle r\rangle\gamma_m)\,(\alpha g_m + k_{mat} + \alpha\gamma_m\langle r\rangle)} \\
&= \frac{g_m\left[k_s P^b + k_s^0(1 - P^b)\right]}{(g_m + \langle r\rangle\gamma_m)}
\end{aligned} \tag{S 7.5}
$$

Substituting S 7.5 into Eq. S 7.2 we get

$$\begin{aligned}\langle r\rangle &= \frac{1}{g_r}\left[k_r - \left\{k_s P^b + k_s^0(1-P^b)\right\} + \frac{g_m\left[k_s P^b + k_s^0(1-P^b)\right]}{(g_m + \langle r\rangle\gamma_m)}\right] \\ &= \frac{1}{g_r}\left[k_r - \frac{\gamma_m\langle r\rangle\left\{(k_s - k_s^0 P^b + k_s^0)\right\}}{(g_m + \langle r\rangle\gamma_m)}\right] \\ &= \frac{1}{g_r}\left[k_r - \frac{\gamma_m\langle r\rangle\left(k_s k_p k_b\langle r\rangle + g_p k_u k_s^0\right)}{(g_m + \langle r\rangle\gamma_m)\left(k_p k_b\langle r\rangle + g_p k_u\right)}\right] \qquad \text{[using S 4.2]} \qquad \text{(S 7.6)}\end{aligned}$$

Eq. S 7.6 gives a polynomial in $\langle r\rangle$ and solving this we obtain $\langle r\rangle$ as a function of $k_r$ along with parameters $g_r, k_p, g_p, k_u, k_b, k_s^0, k_s, g_m, \gamma_m$.
